# Mutant Huntingtin exon1 protein detected in mouse brain with neoepitope antibody: effects of CAG repeat expansion, MSH3 silencing, and aggregation

**DOI:** 10.1101/2024.12.31.630891

**Authors:** Ellen Sapp, Adel Boudi, Andrew Iwanowicz, Jillian Belgrad, Rachael Miller, Riannon Robertson, Daniel O’Reilly, Ken Yamada, Yunping Deng, Marion Joni, Xueyi Li, Kimberly Kegel-Gleason, Anastasia Khvorova, Anton Reiner, Neil Aronin, Marian DiFiglia

## Abstract

*HTT1a* was identified in human and mouse Huntington’s disease brain as the pathogenic exon 1 mRNA generated from aberrant splicing between exon 1 and 2 of *HTT* that contributes to aggregate formation and neuronal dysfunction.^1^ Detection of the huntingtin exon 1 protein (HTT1a) has been accomplished with fluorescence-based reporter assays (Meso Scale Discovery, Homogeneous Time Resolved Fluorescence) and immunoprecipitation assays in Huntington’s disease knock-in mice but direct detection in homogenates by gel electrophoresis and western blot assay has been lacking. Subcellular fractions prepared from mouse and human Huntington’s disease brain were separated by gel electrophoresis and probed by western blot with neo-epitope monoclonal antibodies 1B12 and 11G2 directed to the C-terminal eight residues of HTT1a. In caudate putamen of an allelic series of 6 month old Huntington’s disease knock-in mice (Q50, Q80, Q111, Q140 and Q175) HTT1a migration was inversely correlated with CAG repeat length and appeared as a SDS soluble high molecular mass smear in Q111, Q140 and Q175 mice but weakly in Q80 and not in WT mice or Q50 indicating a CAG repeat size threshold for detecting HTT1a. HTT1a immunoreactivity diminished if 1B12 and 11G2 antibodies were preincubated with an eight amino acid peptide containing the C-terminus of HTT1a but not with unrelated peptide sequence. Migration of HTT1a and its high molecular mass smear changed with age in caudate putamen of Q111, Q175 and YAC128 mice. Treating Q111 mice with siRNA to *MSH3*, a modifier of CAG repeat expansion, significantly reduced levels of the high molecular mass smear indicating that the effects of curbing CAG repeat expansion were quantifiable. A prominent 56-60 kDa doublet detected by 1B12 and 11G2 antibodies in lysates from human Huntington’s disease brain was not blocked by preincubation with C-terminal HTT1a blocking peptide and also appeared in brains of Parkinson’s disease patients. 1B12 and 11G2 antibodies did not immunoprecipitate HTT proteins from either Huntington’s disease mouse or human brain lysates using conditions that pulled down full length HTT with anti-HTT antibody 2B7. Altogether these data show that 11G2 and 1B12 antibodies can be used in western blot assays to track and quantify immunoreactive HTT1a levels, solubility, and subcellular localization in Huntington’s disease mouse brain.

**Abbreviated Summary:** Sapp et al., report that pathogenic exon 1 protein HTT1a is detected in brain of mouse models of Huntington’s disease by direct western blot assay using monoclonal antibodies 11G2 and 1B12. Lowering *MSH3* mRNA in the caudate putamen to prevent CAG repeat expansion reduced levels of HTT1a.

## Introduction

An mRNA arising from aberrant splicing between exon 1 and 2 in the Q150 Huntington’s disease mouse model generates a polyadenylated mRNA named *HTT1a* that is CAG repeat length dependent and also detected in Huntington’s disease patient brain and fibroblasts.^1,2^ *HTT1a* mRNA was proposed to give rise to the toxic huntingtin exon 1 protein (HTT1a) expressed in the R6/2 transgenic mouse model.^3,4^ R6/2 mice are more severely affected than Huntington’s disease knock-in mice because they exhibit phenotypes at an earlier age including large nuclear aggregates in the caudate putamen, behavioral deficits, and early death at about 16-20 weeks. The MW8 antibody made to the C-terminal eight amino acids of exon 1, AEEPLHRP, has been used as a surrogate marker of HTT1a in immunoprecipitation assays, Meso Scale Discovery (MSD) and Homogeneous Time Resolved Fluorescence (HTRF) assays.^1,5^ Antibody MW8 paired with antibody 4C9 or 2B7 had been used in HTRF assays to detect soluble and aggregated forms respectively of HTT1a.^6,7^ S830 is a sheep polyclonal antibody made to exon 1-53Q and along with MW8 mouse monoclonal antibody was considered the most sensitive for detecting aggregates by immunostaining^8^ and was paired with MW8 for use in immunoprecipitation assays to detect HTT1a in Q150 knock-in mice.^2^ Fluorescence-based assays are sensitive high throughput bioassays that can be costly and labor intensive. Detection of HTT1a by HTRF assay requires considerable mouse brain tissue.^7^ A sensitive and direct detection of HTT1a by sodium dodecyl sulphate-polyacrylamide gel electrophoresis (SDS-PAGE) and western blot would provide a rapid, cost-effective method that utilizes smaller tissue samples.

HTT1a has been identified in Huntington’s disease knock-in mice and YAC128 transgenic mice using HTRF, MSD and immunoprecipitation assays^2,7,9,10^ but direct detection by western blot has been elusive. Monoclonal neo-epitope antibodies (P90 1B12 and 11G2) directed to the C-terminal 83-90 of HTT1a (based on HTT with 23 Qs) recently developed and characterized,^11-13^ that are highly sensitive for detecting purified recombinant HTT1a by western blot compared to other antibodies directed to N-terminal HTT fragments, such as MW8, offered the possibility of detecting HTT1a using SDS-PAGE and western blot. Here we tested the sensitivity of neo-epitope antibodies to identify HTT1a by SDS-PAGE and western blot in brain samples from YAC128 transgenic mouse and Huntington’s disease knock-in mice from an allelic series (Q50, Q80, Q111, Q140 and Q175) and compared the results with R6/2 transgenic Huntington’s disease mice which express human exon 1. In Huntington’s disease knock-in mouse models HTT1a appeared as a band and as a high molecular mass (HMM) smear measurable by densitometry. The migration and solubility of HTT1a in SDS-PAGE depended on CAG repeat length, subcellular compartment, and age of mice. Silencing MutS Homolog 3 (*MSH3*) mRNA which encodes a mismatch repair protein that drives CAG expansion in Q111 mice attenuated HTT1a smear in caudate putamen, suggesting the use of 1B12 and 11G2 antibodies is a sensitive quantifiable readout for HTT1a aggregation.

## Materials and Methods

### Sources of Human and Mouse Brain Tissue

**Supplementary Table 1** lists the human brain tissue, regions, sources, and references to our prior publications where brain tissues were used.^14-16^ All human tissues were de-identified and stored frozen at -80°C and age, sex, and CAG repeat length determined from PCR analysis of brain tissue are noted. Huntington’s disease Vonsattel grade of striatal pathology^17^ is included in the table when available.

Mouse brain lysates prepared fresh and stored frozen from prior studies were used.^18-21^ The animal protocols for these studies were approved by the Massachusetts General Hospital Subcommittee on Research Animal Care (SRAC)-OLAW protocol #2004N000248 and University of Massachusetts Chan Medical School Institution Animal Care and Use Committee (IACUC PROTO202000010). All procedures conform to the United States Department of Agriculture Animal Welfare Act, the ‘Institute for Laboratory Animal Research Guide for the Care and Use of Laboratory Animals’ and followed institutional guidelines. Huntington’s disease knock-in mice (Q50, Q80, Q111, Q140 and Q175) were obtained from Jackson laboratories. These mice have the mouse exon 1 replaced with a mutant version of human exon 1 and are on C57BL/6 background. CAG repeat determined from tail DNA was provided by the supplier at the time of delivery and the range for each Huntington’s disease mouse line is reported in **Table 1**. YAC128 mice are on FVB background. For WT, Q111, Q175 and YAC128 mice, male and female mice were used. For WT, Q50, Q80, Q111, Q140 and Q175 allelic series mice, only male mice were used. The experimenters were not blinded to the genotypes or treatment conditions of the mice. A total of 96 mice were used in this study and no samples were excluded. Mice were euthanized and the brain was removed and fresh-frozen or a cardiac perfusion with PBS was performed before brain removal and freezing at -80°C. The cortex from a 12 week old R6/2 mouse on C57BL/6 background was provided by Dr. Vanita Chopra.^22^ The CAG repeat number of 135 was determined at the MGH Genomics Core by PCR and capillary electrophoresis of PCR products.

**Table 1.** Changes in molecular mass of HTT1a protein in western blots probed with antibody 1B12: Effects of CAG repeat number and gel type in Huntington’s disease (HD) mice with different CAG repeats

| HD Mouse model | CAG repeats | Molecular mass of HTT1a protein |  |
| --- | --- | --- | --- |
|  |  | 3-8% Tris-acetate gel | 12% Bis-tris gel |
| R6/2 | 135 | 85-90 kDa | 60 kDa |
| Q80 | 84 | 65 kDa |  |
| Q111 | 108-119 | 72-80 kDa | 56 kDa |
| Q140 | ~140 | 100 kDa |  |
| Q175 | 185-202 | 130-140 kDa | 90 kDa |
| YAC128 | ~128 | 80 kDa |  |

For analysis of *MSH3* silencing, samples of an equal number of male and female Q111 mouse caudate putamen from a prior study^19^ and from a new cohort of mice that were injected bilaterally (125 µg per ventricle) in the lateral ventricle with di-valent siRNA directed to a non-targeting control (NTC) or *MSH3* (MSH3-1000) at three months and sacrificed at 5 months were used. Sequence and chemistry of the siRNAs was previously published.^19^ Mice with confirmed genotypes were randomly assigned to treatment groups, with males and females balanced. To help control for variability between injection sessions, the mice in each injection session are injected with either NTC or siRNA targeting *MSH3*. Mice were anesthetized with isoflurane and monitored throughout the entire intracerebroventricular injection procedure. The mice were maintained on a heat pack the entire time under anesthesia. When the mice were no longer responsive to a touch pinch, fur from the head was shaved, and the mice were placed on the stereotactic frame. A stereotactic procedure was used to locate the lateral ventricle. In brief the skull was exposed using aseptic technique. Bregma was identified, and the stereotactic coordinates were used to locate the skull region over the location of the lateral ventricle (mediolateral ±1 mm, posterior -0.2 mm, ventral -2.5 mm). A burr hole was drilled at the appropriate medial-lateral position and the 33-gauge needle was positioned at ventral -2.5 mm. The divalent siRNA was infused at a rate of 750 nL/min for a total of 2.5 *μ*l. This procedure was repeated in the other hemisphere. A total volume of 5 *μ*l was delivered to each mouse. After the bilateral injections were completed, the skin incision was sutured, and the mice received a subcutaneous injection of Meloxicam ER. The mice were placed on a heat pack in a recovery cage, monitored until sternal, and then checked daily for the first 72 hours post-procedure and weekly throughout the study. After brains were harvested, they were cut into 1mm coronal blocks. The integrity of the ventricular spaces and the brain surrounding the lateral ventricles at the level of the caudate putamen were checked. The trajectory of the needle track could be followed in some brains. When routine histology was performed the lateral ventricle and adjacent tissue were inspected to verify its integrity. Needle punches of the caudate putamen were taken from brain slices to make crude homogenates (CH, see below).

### Preparation of lysates for SDS-PAGE from human and mouse brains

Lysates from brain tissue were prepared using one of two basic methods reported in our prior studies.^20,23^ Pieces of human putamen or cortex (1-2 mm^3^) or mouse caudate putamen were homogenized in 10mM HEPES pH7.2, 250mM sucrose, 1mM EDTA + Protease inhibitor tablet (Roche) + 1mM NaF + 1mM Na_3_VO_4_. An aliquot of the CH was removed, and, in some cases, the sample was centrifuged at 800xg for 15 minutes at 4°C. The supernatant (S1) from this spin corresponds to the cytoplasmic fraction and the pellet (P1), which was resuspended in the same buffer, includes nuclear proteins. For most experiments reported here the CH was used as indicated in figure legends.

A second method of preparation of brain lysates required a different buffer. In brief, the entire caudate putamen per mouse brain were homogenized in 3 ml of 0.32M sucrose, 10mM DTT + Protease inhibitor tablet (Roche) and 100 µl was removed as a CH. Some lysates that had been prepared in this manner were used in a published study of 6-month-old WT, Q50, Q80, Q111, Q140 and Q175 mice.^20^ The remaining samples that were stored frozen were used in the current study.

Protein extraction using mild detergent was tested by homogenizing 6 month WT and Q111 caudate putamen on ice in 20mM Tris pH7.4, 150mM NaCl, 1% Triton X-100, 0.1% NP40, incubating 15 minutes on ice then centrifuging at 16000xg 15 minutes as 4°C following our previous method.^24^ The supernatant was removed and the pellet was resuspended in the same buffer. 20 µg was separated by SDS-PAGE for western blot analysis.

To solubilize aggregated HTT, two methods were tested including our previously published protocol.^16^ Specifically, 20 µg of CH prepared in 10mM HEPES pH7.2, 250mM sucrose, 1mM EDTA as described above from cortex of R6/2 mice and caudate putamen of 6-month-old WT and Q175 mice were incubated in 100 mM DTT and 8M urea for 30 minutes at room temperature with vortexing every 5 minutes before adding LDS sample buffer (Invitrogen) and separating by SDS-PAGE. A second method described by Landles et al.^10^ uses a sequential treatment with SDS followed by formic acid. 50 µg of R6/2 cortex CH was centrifuged at 13000xg for 15 min at 4°C and the pellet was resuspended in 50 µl 1x SDS buffer (2% SDS, 5% beta-mercaptoethanol, 15% glycerol). Samples were boiled for 10 minutes, sonicated for 20 seconds, centrifuged at 13000xg for 15 min at 4°C and the supernatant was removed as the SDS soluble fraction. 100 µl formic acid was added to the pellets and the samples were incubated at 37°C with shaking at 250 rpm for 1 hour, then dried overnight under vacuum in a desiccator. Dried formic acid pellets were neutralized with 1M Tris pH8.6 before adding LDS sample buffer, boiled for 10 minutes and the entire sample was separated by SDS-PAGE as described below.

### SDS-Page and western blot

Protein concentration was determined using the Bradford assay (Bio-Rad Protein assay). Equal amounts of protein were prepared in 1X LDS buffer (Invitrogen) + 100mM DTT, boiled for 5 minutes, separated by SDS-PAGE using 3-8% Tris-acetate (NuPAGE 15-well, 1.5mm, Invitrogen or Criterion 26-well, 1.0mm, Bio-Rad) with Tricine running buffer (Bio-Rad) or 4-12% Bis-tris gels (Criterion 26-well, 1.0mm, Bio-Rad) or 12% Bis-tris gels (NuPAGE 15-well, 1.0mm, Invitrogen or Criterion 26-well, 1.0mm, Bio-Rad) with MOPS running buffer (Bio-Rad), at 120V until the dye-front reached the bottom of the gel (∼1.5 hours), then gels were soaked in Tris-glycine transfer buffer + 0.1% SDS for 5 minutes before transfer. We compared wet transfers at 100V for 1 hour and 35V overnight to our normal transfer using TransBlot Turbo (Bio-Rad) and did not see an appreciable change in the detection of the high molecular mass protein, so we transferred proteins to nitrocellulose with the TransBlot Turbo apparatus at high molecular weight setting (25V for 10 minutes) throughout this study. Blots were incubated in blocking buffer (5% blotting-grade blocker (Bio-Rad) in Tris-buffered saline + 0.1% Tween-20 (TBST)) for 1 hour at room temperature, then in primary antibody diluted in blocking buffer at 4°C overnight with agitation. For controls using blocking and unrelated peptides, antibodies were diluted to 5 µg/ml in blocking buffer and incubated with agitation with the HTT1a peptide or unrelated peptide at 26µM for 1 hour at room temperature for the human brain or overnight at 4°C for the mouse brain before applying to the nitrocellulose blot.

### Immunoprecipitation assays

Immunoprecipitation assays were performed based on our previously published protocol.^25^ Frozen mouse caudate putamen or pieces of human cortex were homogenized in GAL4/IP buffer (50mM Tris pH7.2, 250mM NaCl, 5mM EDTA, 1%NP40 plus protease inhibitor tablet (Roche), 1mM NaF, 1mM Na_3_VO_4_) and incubated on ice for 15 minutes before centrifugation at 13000xg for 2 minutes at 4°C. The protein concentration of the supernatant was determined using the Bradford assay (Bio-Rad) and 5 tubes of 1 mg protein in 1 ml GAL4/IP buffer were prepared. Protein A/G Sepharose beads (Santa Cruz Biotechnology) were washed and blocked for 4 hours or overnight in PBS+1%BSA. Brain lysates were precleared for 1 hour in blocked Protein A/G beads before adding 5 µg 1B12, 11G2 or anti-HTT antibody 2B7 or normal rabbit or mouse IgG and incubating overnight at 4°C. To each tube, 30 µl pre-blocked Protein A/G Sepharose diluted 1:1 in GAL4/IP buffer was added and incubated for 2 hours with agitation at 4°C. Sepharose beads were then centrifuged at 4000 rpm for 4 minutes and washed 4 times in 750 µl GAL4/IP buffer. After final wash, Sepharose beads were resuspended in 30 µl 2x LDS sample buffer + DTT, boiled for 5 minutes then 25 µl was analyzed by western blot with 1B12 or anti-HTT antibodies.

### Sources of antibodies and peptides used for western blot

Antibodies and dilutions were as follows: anti-HTT antibody Ab1 (aa1-17^26^, 1:2000, rabbit), MW8 (Developmental Studies Hybridoma Bank, University of Iowa, 1:500, mouse), S830 (generous gift from Dr. Gillian Bates, 1:6000, sheep), 2B7 antibody (N-terminal HTT, Coriell Institute for Medical Research, 1:750, mouse), P90 antibodies (neo-epitope to aa83-90, clones 1B12 and 11G2, Coriell Institute for Medical Research, 5 µg/ml, rabbit), MSH3 (Santa Cruz BioTech #sc-271079, 1:500, mouse), GAPDH (MilliporeSigma #MAB374, 1:10000, mouse), B-actin (MilliporeSigma #A5441, 1:5000, mouse). Blots were washed in TBST, then incubated in peroxidase-labeled secondary antibodies diluted 1:2500 for rabbit IgG and 1:5000 for mouse and sheep IgG in blocking buffer for 1 hour at room temperature. Immunoprecipitation blots were incubated in rabbit or mouse IgG, light-chain specific (Cell Signaling), diluted 1:2000 in blocking buffer. Blots were washed in TBST, and bands were visualized using SuperSignal West Pico PLUS Chemiluminescent substrate (Thermo Scientific) and ChemiDoc XRS+ with Image Lab software (Bio-Rad). Some blots were stripped for 30 minutes with Restore stripping buffer (Thermo Scientific) before washing, blocking and reprobing with a different antibody. Peptides were obtained from ThermoFisher. The sequences were AEEPLHRP for HTT1a C-terminus and YHRLLTCLRNVHKVTTC for the unrelated peptide.

### Pixel intensity quantification

Image analysis was performed using ImageJ software. For HTT, MSH3 and loading controls, bands were manually circled, and the area and average intensity were determined. The total signal intensity of each band was calculated by multiplying the area by the average intensity. For HTT1a levels in YAC128, allelic series and Q111 mice treated with *MSH3* siRNA, the average intensity of the smear and band was measured in equal areas for each lane. The dashed lines in one lane of western blots represent the areas measured in all lanes. For the allelic series, the average signal intensity of the smear obtained in the WT lanes was subtracted as background from the average signal intensity of the smear in the Q80, Q111, Q140 and Q175. The signal intensities were normalized to the loading control signal of B-actin or GAPDH.

### Neuronal differentiation

Experiments were performed with oversight of Human Embryonic Stem Cell Research Oversight Committee (ESCRO Committee) through the MassGeneral Brigham Institutional Biosafety Committee (PIBC) (ESCRO#: 2015-01-02 and PIBC Reg# 2017B000023) using human induced pluripotent stem cells (iPSCs) CS09iCTR-109n4 (CS vial ID: 1034860) and CS09iCTR-109n5 (CS vial ID: 1034589) hereafter referred as HD109 (clone n4 and n5) described by Mattis et al., 2015.^28^ iPSCs were differentiated into cerebral cortex neurons as described^29,30^ with modifications. Briefly, neural induction occurred on confluent iPSCs plated on Matrigel (Corning, 354277) for 10 days in Neural Induction Medium. Neuroepithelial sheets were lifted with dispase (Stemcell technologies, catalog 07923) and plated on Matrigel. FGF treatment (20ng/mL, Gibco, PHG0023) in Neural Maintenance Medium (NMM) was initiated on day 14 and stopped at Day 18 where neural rosettes were observed. Upon successful neural development, cells were cultured continuously in NMM with BDNF (10ng/mL, Gibco, PHC7074) and GDNF (10ng/mL, Gibco, PHC7075) 42 days after the start of the neural induction along with anti-mitotic inhibitors 5-Fluoro-2’-Deoxyuridine (1µM, Sigma-Aldrich, F0503) and Udirine (1µM, Sigma-Aldrich, U3750). Neuron cultures were characterized using an antibody panel was performed by western blot and immunocytochemistry to endorse the proper neuronal differentiation. Neurons were harvested in 10mM HEPES pH7.2, 250mM sucrose, 1mM EDTA + PI tablet (Roche) + 1mM NaF + 1mM Na_3_VO_4_ and homogenized on ice before determining protein concentration using the Bradford method (Bio-Rad).

### Statistical analysis

Statistical analyses were performed using GraphPad Prism v10.4.1 software. Data were tested for Gaussian distribution using the Shapiro-Wilkes test. Those data sets with normal distribution were subject to unpaired t tests or one-way ANOVA with Tukey’s multiple comparison test and the data set that was not normally distributed was subject to a Mann Whitney test. Pearson r test was used for correlation analyses. Statistical test, sample size and p values are listed in the figure legends.

## Results

### Detection and solubility of HTT1a protein in brain of Huntington’s disease knock-in mice and YAC128 transgenic mice is dependent on age and CAG repeat length

We examined lysates from Q111 Huntington’s disease mice which belong to an allelic series that have mouse exon 1 replaced with a mutant version of human exon 1 and one R6/2 mouse that has a human exon 1 transgene (**Fig. 1**). Using 3-8% Tris-acetate gradient gels for protein separation, 1B12 and 11G2 antibodies detected immunoreactive HTT1a by western blot at around 72 kDa in soluble (S1) fractions prepared from caudate putamen of two Q111 Huntington’s disease knock in mice and around 80 kDa in one R6/2 mouse cortex. These bands were absent in Q111 samples or significantly diminished in R6/2 lysates by preincubation with a C-terminal HTT1a peptide (**Fig. 1A, B**). A broad HMM smear was also detected in the R6/2 lysates by 1B12 and 11G2 antibodies and markedly reduced by antibody preincubation with C-terminal HTT1a peptide. Preincubation with an unrelated peptide had no effect (**Fig. 1A, B**). HTT1a bands migrated faster if 12% Bis-tris gels were used for protein separation (see **Table 1** for molecular mass comparisons) and the smear in R6/2 was more compressed at the top of the blot (**Supplementary Fig. 1**). These results suggest the immunoreactive bands and smear detected by 1B12 and 11G2 recognized HTT1a. To further support specificity, immunoprecipitation assays were performed. Antibodies 11G2 and 1B12 did not immunoprecipitate HTT1a in brain lysates from Huntington’s disease Q140 or Q111 mice whereas under the same conditions full length WT and mutant HTT were immunoprecipitated by antibody 2B7 (**Supplementary Fig. 2).** To determine if HTT1a detection in brains of Huntington’s disease knock-in mice could be improved by methods that solubilize aggregates,^10^ lysates were treated with urea, or SDS and formic acid, or by using a gentler extraction in Tris buffer with Triton X-100 and NP40. None of these preparations of the lysates improved detection of HTT1a (**Supplementary Fig. 3A-D**).

**Fig. 1.**
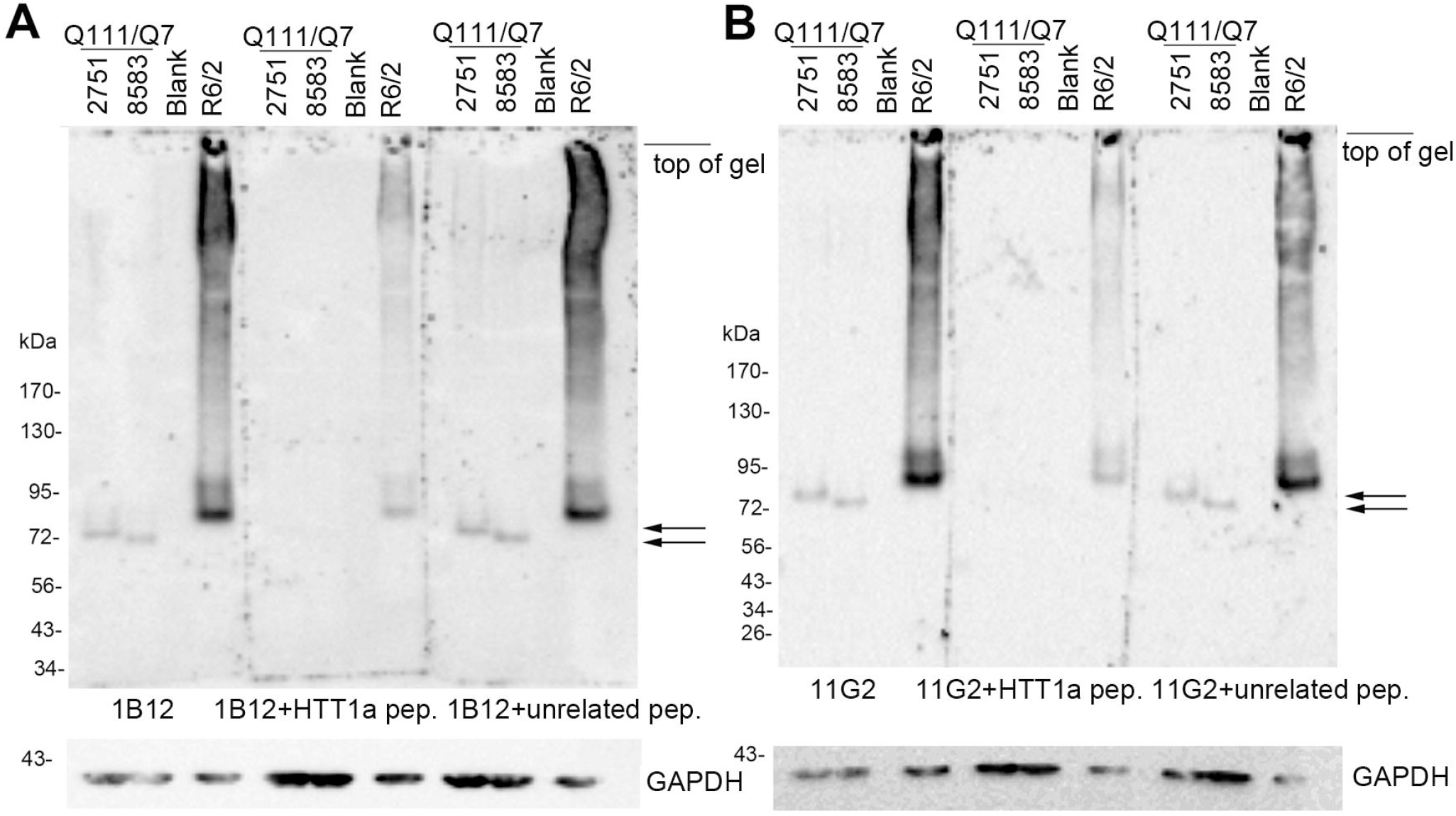
Peptide blocking of HTT1a antibodies 1B12 and 11G2. The signal at 72-75kDa (at level of arrows) in Huntington’s disease Q111 mice is absent when antibodies 1B12 (**A**, middle blot) and 11G2 (**B**, middle blot) are preincubated with the HTT1a C-terminal peptide AEEPLHRP before western blot analysis. The intensity of the HTT1a bands and smear in the R6/2 cortex is greatly reduced by antibody preincubation with the blocking peptide. HTT1a detection by 1B12 (**A**) and 11G2 (**B**) antibodies is unaltered by preincubation with an unrelated peptide (compare left and right blots in **A** and **B**). 20 µg of Q111 caudate putamen S1 fraction samples 2751 (118 CAG repeats) and 8583 (113 CAG repeats) and R6/2 cortex CH (crude homogenate) were separated on 3-8% Tris-acetate 15 well gels. At the top of the blots, mouse numbers are positioned at the center of each lane. “Blank” marks a lane without protein.

Next the effects of age and subcellular compartment on HTT1a expression were explored in different Huntington’s disease mouse models. The presence of HTT1a in crude homogenates, S1, and P1 fractions of caudate putamen from 6-month-old Q111 mice were analyzed together by western blot (**Fig. 2A**). HTT1a migrated as a discrete band to about 75 kDa in S1 fractions and as a HMM smear in crude homogenates and P1 fractions (**Fig. 2A**). In S1 fractions from 6-, 12- and 24-week-old Q111 mice compared in the same western blot, HTT1a migrated to about 75 kDa (**Fig. 2B, arrow**), but levels diminished with age to about 50% at 24 weeks when HTT1a solubility was replaced by a HMM smear (**Fig. 2B**). No smear appeared in WT mice lysates at 24 weeks. HTT1a was also examined in Q175 mice of different ages where it migrated to about 130 kDa (**Fig. 2C**, arrow) and decreased in intensity with age between 2 months and 10 months coinciding with a marked rise in the SDS soluble smear (**Fig. 2C**). HTT1a smear was also significantly increased at 8 months compared to 4 months in lysates from YAC128 mice (**Fig. 2D, Supplementary Fig. 3E**). Altogether these data show that HTT1a migration by western blot was age-dependent migrating more slowly by SDS-PAGE with increased CAG repeat expansion, although other factors affecting HTT1a migration cannot be ruled out.

**Fig. 2.**
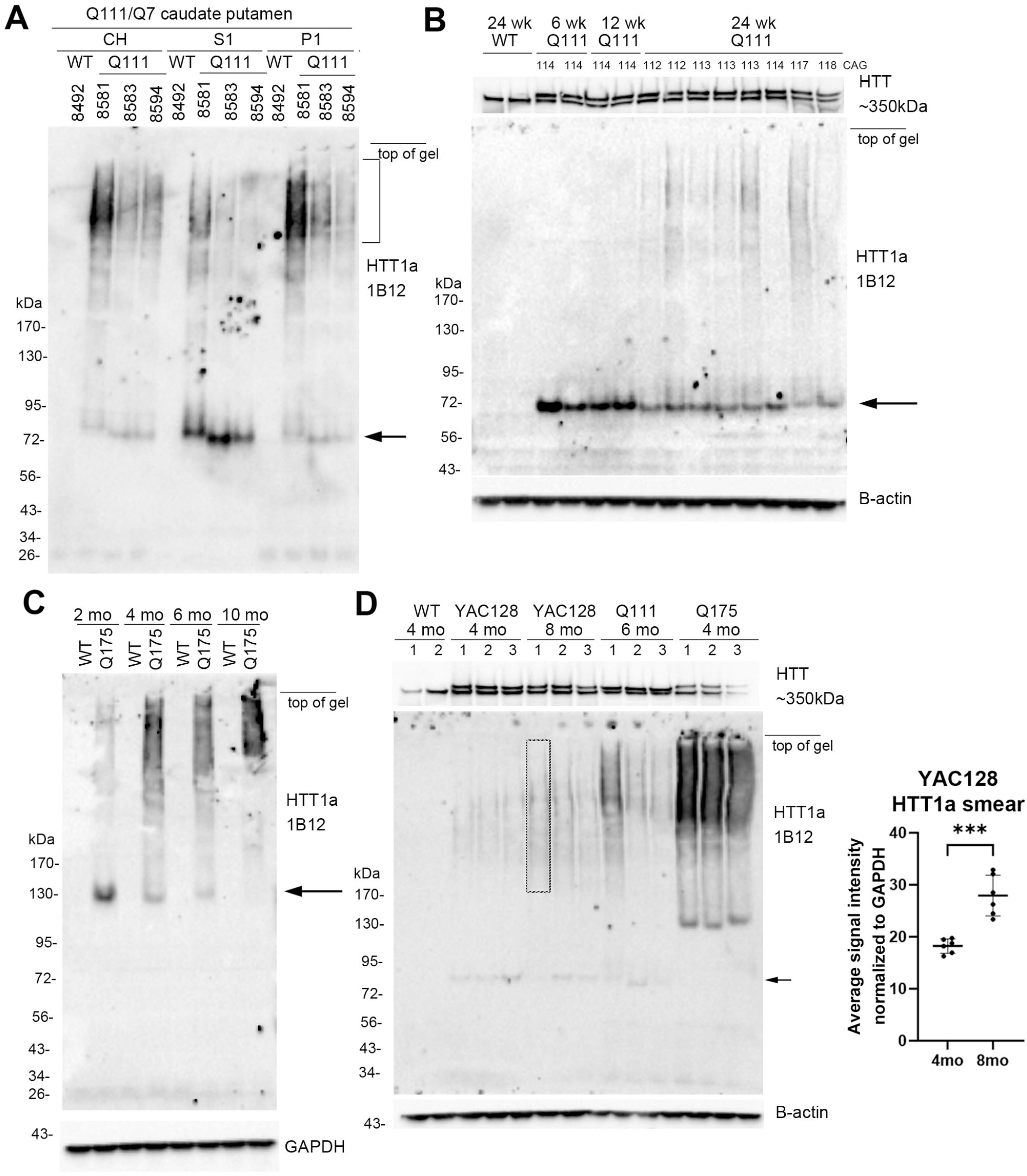
Effects of age on HTT1a expression in Q111, Q175 and YAC128 mice. Changes in levels, molecular mass and SDS solubility. **A.** Crude homogenates (CH) as well as S1 and P1 fractions from the same mice were separated on a 3-8% Tris-acetate 26-well gel and probed with P90 antibody 1B12. HTT1a migrates to 75 kDa (arrow) and is most expressed in the S1 fraction whereas it appears as a high molecular mass (HMM) smear (bracket) in CH and P1 fractions. **B.** Western blot of S1 fractions of caudate putamen from WT mouse at 24 weeks and Q111 mice at 6, 12 and 24 weeks. 20 µg samples were separated on a 15-well 3-8% Tris-acetate gel. HTT1a migrates to about 75 kDa (arrow), and intensity decreases progressively between 6 weeks and 24 weeks. Note that at 24 weeks HTT1a smear appears in Q111 mice but not in WT mouse. CAG repeat was determined from tail DNA and is indicated at the top of the blot. Blot was re-probed with anti-HTT antibody Ab1 to detect full-length HTT. **C.** Western blot of crude homogenates of WT and Q175 mice at 2, 4, 6 and 10 months. 20 µg samples were separated on a 3-8% Tris-acetate 26-well gel. HTT1a band migrates to about 130 kDa (arrow), and intensity decreases with increased age from 2 months to 10 months. In contrast the presence of HMM HTT1a smear increases with age. HTT1a is not detected in the WT mice. **D**. Western blot of CH prepared from caudate putamen of 4- and 8-month-old YAC128 mice show a significant increase in the HTT1a smear from 4 to 8 months (***p=0.0002, t=5.692, df=10, two-tailed unpaired t test, n=6 per group, each dot represents one mouse and bars are mean±SD). Three of the six samples are shown in this figure. Images for analysis of n=6 samples per group are shown in **Supplementary Fig. 3E**. The HTT1a band in the YAC128 mice runs at about 80 kDa (small arrow) which is slightly larger than in the Q111 mice, as expected. Note that the loading controls, which are B-actin for **B** and **D** and GAPDH for **C**, show equal signal intensities indicating equal protein loading. At the top of the blots, mouse numbers or CAG repeats are positioned at the center of each lane. Full blots are shown in **Supplementary Fig. 5**.

We also compared HTT1a expression in Huntington’s disease knock-in mice that had different CAG repeats in exon 1 (**Fig. 3A**). CH from caudate putamen of 6-month-old WT, Q50, Q80, Q111, Q140 and Q175 mice showed differences in HTT1a migration when proteins were separated in 3-8% Tris-acetate gradient gels. HTT1a was ∼65 kDa in Q80 mice, 75 kDa in Q111 mice, 100 kDa in Q140 mice and 130 kDa in Q175 mice (**Fig. 3A** small arrows, **Table 1**). The intensity of HTT1a also significantly increased with CAG repeat size (graphs in **Fig. 3A**). A HTT1a HMM smear appeared in Q111, Q140 and Q175 mice and increased in signal intensity with increase in CAG repeat number (graphs in **Fig. 3A**). HTT1a was not detected in WT or Q50 mice. N-terminal HTT1-17 antibody Ab1 detected full length WT and mutant HTT at about 350 kDa in all mice. In Q175 mice full-length HTT levels were significantly lower suggesting that the marked HTT1a accumulation seen as a smear on western blot in these mice may affect levels of full length HTT (**Fig. 3A,** top).

We wondered if the smear seen with 1B12 and 11G2 was also detected by other anti-HTT antibodies. S830 is a polyclonal sheep antibody raised against exon 1 with 53Q and MW8 is a monoclonal antibody made against exon 1 with 67Q. Both antibodies label aggregates by immunohistochemistry and MW8 and has been used in HTRF based aggregation assays.^6,32^ All three antibodies detected a HMM smear in the R6/2 transgenic mice but only 1B12 detected a smear in samples from Huntington’s disease knock-in mice (**Fig. 3 B, C, D**). This suggests 1B12 antibody is sensitive to a unique polyglutamine conformation in HTT.

**Fig. 3.**
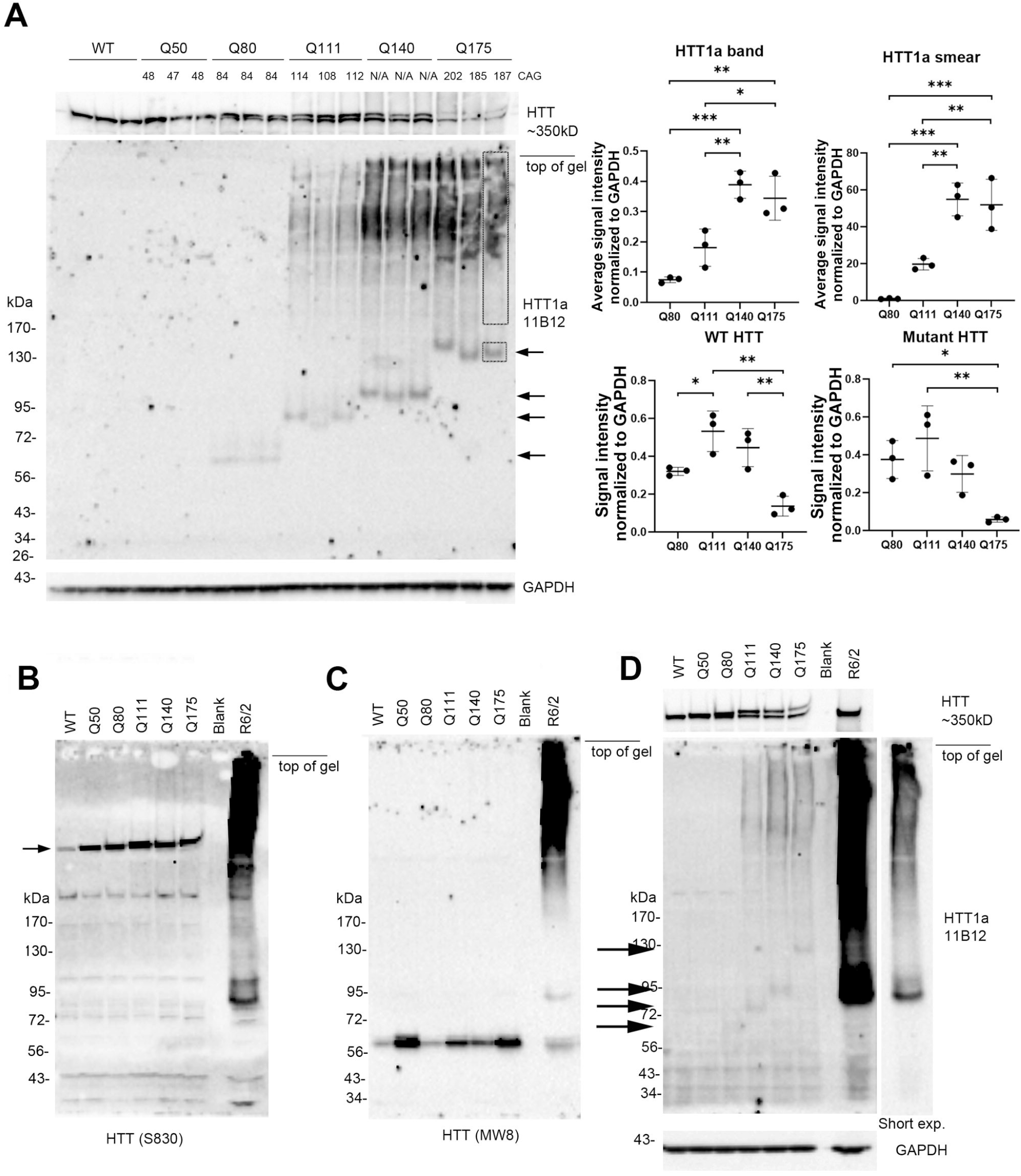
Intensity, size, and solubility of HTT1a in caudate putamen of 6-month-old Huntington’s disease knock-in mice allelic series. **A.** SDS-PAGE and western blot probed with antibody 1B12 in crude homogenates from caudate putamen of Q50, Q80, Q111, Q140 and Q175 mice. 15 µg samples were separated on 3-8% Tris-acetate 26-well gels. Numbers at the top are CAG repeats determined from tail DNA. Large blot shows HTT1a probed with 1B12 and the strip below is loading control for GAPDH. Migration of HTT1a (arrows) slows with increasing CAG repeat from about 65 kDa in Q80 to about 130 kDa in Q175 mice. HTT1a is not visible in WT or Q50 mice. HTT1a smear is seen in lysates from Q111, Q140 and Q175 mice. Strip above large blot is a reprobe with anti-HTT antibody Ab1 and shows WT and mutant HTT. Note decline in levels of full-length WT and mutant HTT in Q175 samples. Graphs at top right show average intensity of HTT1a and HTT1a smear normalized to GAPDH. Boxes in right lane of blot indicate the same areas that were measured in all lanes. The signal measured in WT lanes was subtracted out as background and the signal for non-specific dots in the 3^rd^ lane of Q140 mice and 2^nd^ and 3^rd^ lane of Q175 mice were also subtracted. Bottom graphs show levels of WT and mutant HTT normalized to GAPDH (one-way ANOVA with Tukey’s multiple comparison test, HTT1a band: F=22.75, p=0.0003; HTT1a smear: F=29.04, p=0.00001; WT HTT: F=14.30, p=0.0014; Mutant HTT: F=89.029, p=0.0085; *p<0.05, **p<0.01, ***p<0.01, n=3 per group, each dot represents one mouse and bars are mean ± SD). **B-D**. Comparison of protein detection by antibodies S830, MW8 and 1B12. 15 µg crude homogenates from n=1 each WT, Q50, Q80, Q111, Q140, Q175 striatum and R6/2 cortex were separated on two 3-8% Tris-acetate 15-well gels, one probed with anti-HTT antibody S830 (**B**) and the other with anti-HTT antibody MW8 (**C**). The blot in **C** was stripped and re-probed with 1B12 antibody (**D**). In **B, C,** and **D**, all 3 antibodies detect 1-2 bands in R6/2 mouse cortex at ∼90 and 100 kDa and a prominent high molecular mass SDS soluble smear. S830 detects full length HTT (arrow on left) in all cortex samples with a stronger signal for mutant than WT HTT. In **C**, MW8 detects an unknown band at ∼60 kDa in WT and Huntington’s disease knock-in mouse models. In **D**, 1B12 antibody detects HTT1a which migrates in relation to its CAG repeat: at 65 kDa for Q80, 75 kDa in Q111, 100 kDa in Q140 and 130 kDa in Q175 (arrows on left). No CAG length dependent bands are detected in Huntington’s disease knock-in mice with S830 or MW8 (**B, C**). At the top of the blots, mouse labels or CAG repeats are positioned at the center of each lane. “Blank” marks a lane without protein. Full blots are shown in **Supplementary Fig. 6**.

### The HTT1a smear decreases in caudate putamen of Q111 mice after lowering levels of MSH3, a mismatch repair protein

Results above in Huntington’s disease knock-in mice showed that detection of HTT1a by western blot was dependent on its subcellular compartment, age of mice, and CAG repeat length. To determine if HTT1a detected with 1B12 antibody could be quantified by western blot assay, we compared different protein concentrations (5, 10, 20, and 40 µg) from the same Q111 mouse brain and performed densitometry. Results showed a concentration dependent increase in signal for HTT1a, signifying the antibody was sufficiently sensitive to use for a quantitative assay (**Fig. 4A**).

**Fig. 4.**
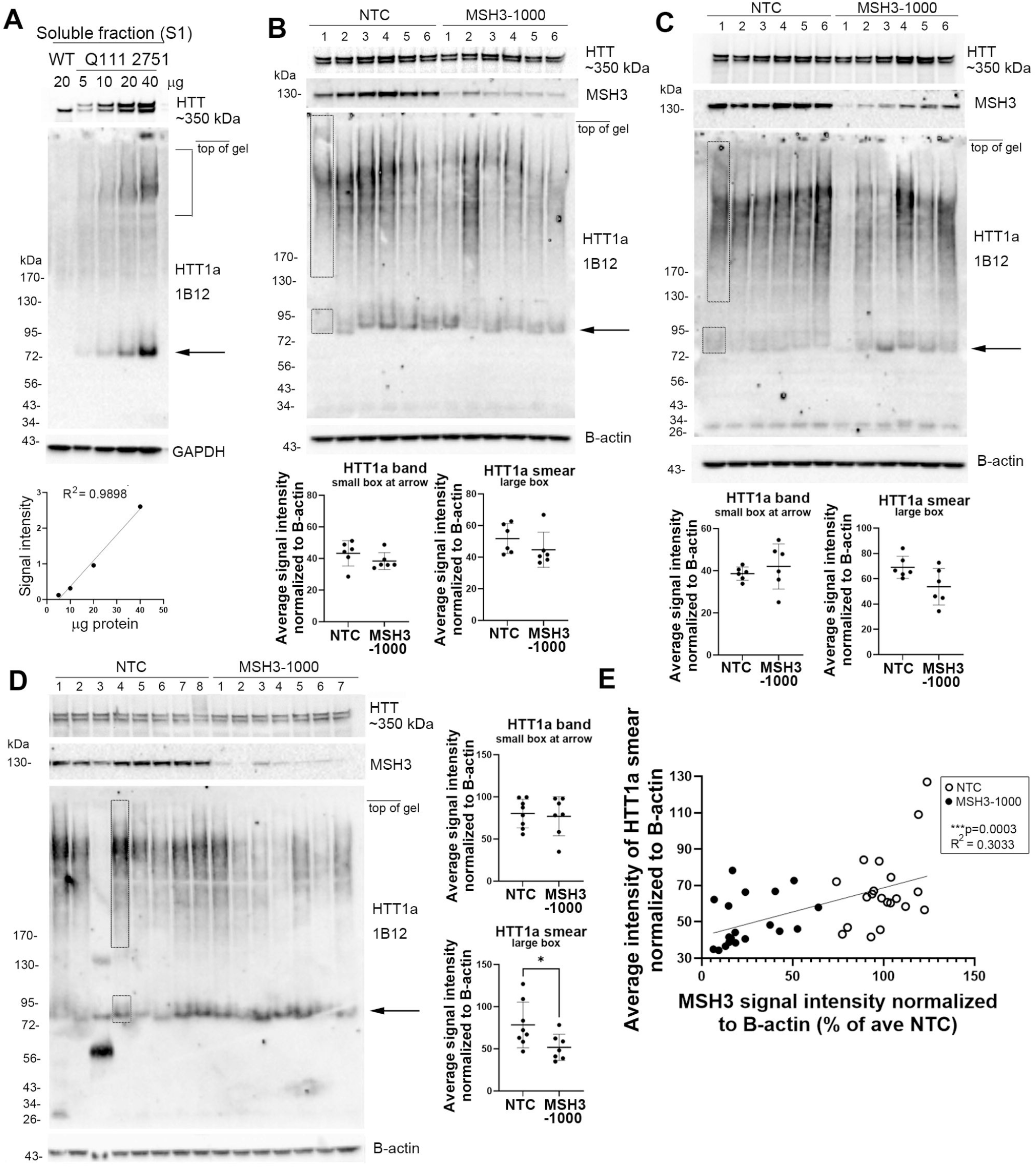
Levels of HTT1a in Huntington’s disease mice caudate putamen: Effects of protein concentration and treatment with siRNA to *MSH3*. **A.** Signal for HTT1a detected with 1B12 antibody is concentration dependent. S1 fraction from Q111 mouse 2751 was loaded at different protein concentrations for SDS-PAGE (5-40 µg) using a 3-8% Tris-acetate 15-well gel. Western blot probed with 1B12 shows signal for 75 kDa band (arrow) increases with increasing protein concentration (graph below, Pearson’s r test, r=0.9949, R^2^ =0.9898, p=0.0051 (two-tailed)). HTT1a smear at bracket is also concentration dependent. Similarly full length HTT detected with Ab1 (above) and GAPDH (below) increase with protein concentration. The 75 kDa band at level of arrow was detected in all Q111 samples but not WT. **B, C, D.** Effects of reducing levels of *MSH3* mRNA in Q111 mice on HTT1a levels. Three experiments show that HTT1a smear is reduced in most lanes where MSH3 levels are markedly reduced. Graphs show intensity of HTT1a smear and 75 kDa band (arrow in **B, C and D**) where each dot represents one mouse. Equal areas of the smear and 75 kDa band as indicated by dashed lines in the first lane in **B** and **C** and fourth lane in **D** were measured by densitometry. Graphs show signal intensity of 75 kDa band is not changed but HTT1a smear is significantly reduced in some lysates from siRNA *MSH3*-treated mice compared to NTC (**D**, *p=0.0395, t=2.289, df=13, two-tailed unpaired t test, N=8 NTC and 7 MSH3). Blots from the same samples were also probed with anti-HTT antibody Ab1 (aa1-17), anti-MSH3 and anti-B-actin antibodies. **E.** There is a significant correlation between MSH3 and HTT1a signal intensities when the data from the three experiments were combined (Pearson’s r test, r=0.5507, R^2^ = 0.3033; ***p=0.0003 (two-tailed), n= 20 NTC and 19 MSH3 mice, each dot represents one mouse). 20 µg crude homogenate samples were separated on 3-8% Tris-acetate 15-well gels in **B** and **C** and a 26-well gel in **D**. At the top of the blots, mouse numbers are positioned at the center of each lane. Full blots are shown in **Supplementary Fig. 7**.

The mismatch repair protein MSH3 is required for CAG repeat expansion and nuclear localization of mutant HTT.^31^ Previously we showed that treating 3-month old Q111 mice with MSH3-1000, a divalent siRNA targeting *MSH3*, lowered MSH3 protein by 54% and inhibited CAG repeat expansion at 5 months compared to treatment with a non-targeting control (NTC).^19^ In lysates from caudate putamen of mice treated with MSH3-1000 siRNA from this study along with two other cohorts (N=6-7 mice/group/study), there was a significant depletion in levels of HTT1a smear that positively correlated with reduction in MSH3 protein levels (**Fig. 4B-E**, R^2^=0.3033, p=0.0003, n=19-20 mice per group from three studies). HTT1a smear was conspicuously lowered in mice where MSH3 protein was reduced by 60-75% (**Fig. 4B-D**). In the same mice, there was no change in levels of full-length HTT. These results demonstrate that the changes in HTT1a smear detected by 1B12 results from CAG repeat expansion in *HTT1a* and can be used to quantify the effects of treating Q111 mice with agents that inhibit somatic expansion. Full blots for mouse data are shown in **Supplementary Figs. 5-9**.

### 1B12 and 11G2 do not detect immunoreactive HTT1a by western blot in lysates from Huntington’s disease brain and human HD109Q neurons

*HTT1a* mRNA has been detected in some samples from human Huntington’s disease postmortem brain and patient fibroblasts.^2^ Therefore, we applied similar methods of analysis used in Huntington’s disease mice to examine Huntington’s disease postmortem brain. In S1 and CH fractions that were separated in 3-8% Tris-acetate (**Supplementary Fig. 4A,B**) or 12% Bis-tris (**Supplementary Fig. 4C**) gels, 1B12 antibody detected a doublet at 56-60 kDa that was more prevalent in the human Huntington’s disease putamen than in control brain. These bands were not blocked when 1B12 antibody was preincubated with the C-terminal HTT1a blocking peptide (**Supplementary Fig. 4A**). The 56-60 kDa bands also appeared in post-mortem cortex from three Parkinson’s disease patients (**Supplementary Fig. 4C**) suggesting that the non-specific band may be elevated in disease brain. Immunoprecipitation with 1B12 or 11G2 from human Huntington’s disease brain lysates of adult-onset HD patients with 42 and 53 CAG repeats in the *HTT* allele and re-probing with 2B7 antibody failed to detect HTT immunoreactive proteins. However, using similar assay conditions, proteins immunoprecipitated with 2B7 antibody expressed full length WT and mutant HTT when detected with Ab1 antibody (**Supplementary Fig. 4D-F**). Finally, we examined lysates from iPSC derived medium spiny neurons and cortical neurons in WA09 and HD109Q lines. Despite the higher CAG repeat in the HD109Q line, only the 56 kDa band was detected in both control WA09 and HD109Q lines (**Supplementary Fig. 4G).** Altogether these findings suggested that unlike Huntington’s disease mouse brain, 1B12 and 11G2 antibodies did not detect HTT1a in human Huntington’s disease brain or in iPSC derived human HD109Q neurons by western blot assays. Full blots for human data are shown in **Supplementary Fig. 10**.

## Discussion

Aberrant splicing between exon 1 and exon 2 of *HTT* due to the presence of a cryptic polyadenylation in intron 1 produces an exon 1 mRNA named *HTT1a* and encoded protein HTT1a that terminates at a proline residue (aa 90, based on 23 CAGs).^1,2^ This fragment was identified as translating the endogenously generated HTT exon 1 protein product that may nucleate and recruit aggregates. *HTT1a* mRNA has been detected in human Huntington’s disease brain, in a wide range of human peripheral tissues and in Huntington’s disease knock-in mouse models that express human exon 1 and its levels of expression are CAG repeat length dependent.^2,9,34^ HTT1a protein levels in human and mouse Huntington’s disease brain have been assessed using immunoprecipitation and HTRF assays with N-terminal HTT antibodies that individually are not specific for the endogenous protein.^2^ Here we showed that direct western blot assays can be used to detect HTT1a in lysates from Huntington’s disease transgenic (R6/2, YAC128) and Huntington’s disease knock-in mouse brain using neo-epitope specific monoclonal antibodies P90-1B12 and 11G2 which were directed to the C-terminal eight amino acids of HTT exon 1 protein.^11,12^ Six-month-old mice from the allelic series of Huntington’s disease knock-in mouse models resolved HTT1a as discrete bands and as prominent HMM smears that varied in migration and signal intensity in relation to CAG repeat length and age of the mice. The smears appeared in 6-month old Q111, Q140 and Q175 mice, consistent with the presence of aggregates that are prevalent by this age with immunostaining^6,35-37^ and in 4- and 8-month old YAC128 mouse striatum concurrent with loss of striatal volume at 3 months of age^38^ and appearance of EM48-positive nuclear aggregates by immunostaining at 2 months.^39^ Moreover, densitometry showed that the HTT1a smear was significantly reduced in caudate putamen of 5-month-old Q111 mice treated at 3 months with siRNA targeting *MSH3*. O’Reilly and colleagues^19^ showed there was no difference in the instability index between the baseline 3-month mice and those treated with *MSH3* siRNA for 2 months whereas mice treated with NTC had a significantly higher instability index suggesting that further expansion was blocked by *MSH3* siRNA. These findings support the presence of HTT1a in mouse Huntington’s disease brain and the feasibility of using western blot detection of HTT1a in Huntington’s disease mice as a biomarker to test the effects of agents that limit CAG repeat expansion.

The HTT1a smear seen in Huntington’s disease mice with 1B12 and 11G2 antibodies may partly correspond to SDS insoluble products identified in Q175 and R6/2 mice using other biochemical assays. HMM products migrating above 300 kDa were found in nuclear fractions from R6/2 mouse brain by agarose gel electrophoresis for resolving aggregates (AGERA) and probed with antibodies 4C9 and MW8.^5^ Also, in nuclear fractions from brain lysates of R6/2 mice SDS insoluble products remained at the top of a western blot probed with S830 antibody.^5^ Consistent with these protein studies, a screen of antibodies by Bayram-Weston et al.^8^ showed that MW8 and S830 antibodies were the most sensitive for identifying nuclear inclusions and diffuse staining in Huntington’s disease transgenic and knock-in mice. The decline in HTT1a smear observed by western blot after lowering *MSH3* mRNA contrasts with the limited effects on aggregate formation seen by immunostaining after silencing *MSH3* mRNA in Q111 mice and Q175 mice.^42,43^ This suggests the HTT1a smear detected by 1B12 and 11G2 in western blot is a fraction of the aggregated mutant HTT present in tissue.

Factors that could explain why 1B12 and 11G2 antibodies did not detect HTT1a in the human Huntington’s disease brain are that protein levels are too low due to marked neuronal loss and/or the protein is sequestered into a nuclear compartment that is not accessible. Also, based on our findings in Huntington’s disease knock-in mice, the CAG repeat size may need to reach 80 to detect the HTT1a band and at least 100 to see a smear. Moreover, immunoprecipitation of HTT1a with the 1B12 and 11G2 antibodies from lysates of human as well as mouse Huntington’s disease brain was unsuccessful using assay conditions that pull down and enrich for full length HTT with an anti-HTT N-terminal antibody. The marked loss of neurons in Huntington’s disease patient neostriatum and cortex compared to Huntington’s disease knock-in mouse and the proliferation of glial cells which are reported to have less somatic instability than neurons^44,45^ could limit levels of HTT1a in human Huntington’s disease brain. Also, RNA sequencing analysis of human Huntington’s disease neostriatum suggests that very large somatic expansions that may be required to detect HTT1a are infrequent and short-lived.^46^ In studies of cortex using single serial fluorescence-activated nuclear sorting (sFANS) and single nucleus RNA sequencing, layer 5a pyramidal neurons in Huntington’s disease postmortem cortex exhibit CAG repeat expansions.^46,47^ Since these neurons were reported as lost early in disease, they may be sparse in our postmortem samples from the cortex.

In summary, our results showed that monoclonal 1B12 and 11G2 antibodies are useful for direct western blot detection of HTT1a in subcellular fractions from brain of Huntington’s disease knock in mice. The sizes and solubility of HTT1a varied with subcellular compartment, age and CAG repeat expansion and these features could be easily compared. Relatively small amounts of tissue are required for detection of HTT1a by SDS-PAGE and western blot, allowing tissues for other assays to be collected from the same mouse brain. Moreover, the ability to quantify changes in levels of HTT1a smear in western blots after inhibiting CAG repeat expansion will be a useful readout for preclinical studies.

## Supplementary material

PDF with 1 supplementary table and 10 supplementary figures is included.

## Data availability

The authors confirm that the data supporting the findings of this study are available within the article and its supplementary material. Raw data is available upon request from the corresponding author.

## References

1. Sathasivam K, Neueder A, Gipson TA, et al. Aberrant splicing of HTT generates the pathogenic exon 1 protein in Huntington disease. Proc Natl Acad Sci U S A. Feb 5 2013;110(6):2366–70. doi:10.1073/pnas.1221891110

2. Neueder A, Landles C, Ghosh R, et al. The pathogenic exon 1 HTT protein is produced by incomplete splicing in Huntington’s disease patients. Sci Rep. May 2 2017;7(1):1307. doi:10.1038/s41598-017-01510-z

3. Mangiarini L, Sathasivam K, Seller M, et al. Exon 1 of the HD gene with an expanded CAG repeat is sufficient to cause a progressive neurological phenotype in transgenic mice. Cell. Nov 1 1996;87(3):493–506. doi:10.1016/s0092-8674(00)81369-0

4. Davies SW, Turmaine M, Cozens BA, et al. Formation of neuronal intranuclear inclusions underlies the neurological dysfunction in mice transgenic for the HD mutation. Cell. Aug 8 1997;90(3):537–48. doi:10.1016/s0092-8674(00)80513-9

5. Landles C, Milton RE, Ali N, et al. Subcellular Localization And Formation Of Huntingtin Aggregates Correlates With Symptom Onset And Progression In A Huntington’S Disease Model. Brain Commun. 2020;2(2):fcaa066. doi:10.1093/braincomms/fcaa066

6. Smith EJ, Sathasivam K, Landles C, et al. Early detection of exon 1 huntingtin aggregation in zQ175 brains by molecular and histological approaches. Brain Commun. 2023;5(1):fcad010. doi:10.1093/braincomms/fcad010

7. Landles C, Osborne GF, Phillips J, et al. Mutant huntingtin protein decreases with CAG repeat expansion: implications for therapeutics and bioassays. Brain Commun. 2024;6(6):fcae410. doi:10.1093/braincomms/fcae410

8. Bayram-Weston Z, Jones L, Dunnett SB, Brooks SP. Comparison of mHTT Antibodies in Huntington’s Disease Mouse Models Reveal Specific Binding Profiles and Steady-State Ubiquitin Levels with Disease Development. PLoS One. 2016;11(5):e0155834. doi:10.1371/journal.pone.0155834

9. Fienko S, Landles C, Sathasivam K, et al. Alternative processing of human HTT mRNA with implications for Huntington’s disease therapeutics. Brain. Dec 19 2022;145(12):4409–4424. doi:10.1093/brain/awac241

10. Landles C, Sathasivam K, Weiss A, et al. Proteolysis of mutant huntingtin produces an exon 1 fragment that accumulates as an aggregated protein in neuronal nuclei in Huntington disease. J Biol Chem. Mar 19 2010;285(12):8808–23. doi:10.1074/jbc.M109.075028

11. Baldo B, Peladan J, Albers J, et al. Development of a HTT exon1-selective MSD immunoassay with novel HTT P90 neo-epitope specific antibodies. 19th Annual Huntington’s Disease Therapeutics Conference; 2024.

12. Missineo A, Tomei L, Alaimo N, et al. Highly selective monoclonal antibodies targeting the HTT exon1 neo-epitope. 19th Annual Huntington’s Disease Therapeutics Conference; 2024.

13. Baldo B, Peladan J, Albers J, et al. Development of an HTT exon1-selective MSD immunoassy with novel HTT P90 neo-epitope-specific antibodies. J Neurol Neurosurg Psychiatry. 2024;95:A32.

14. Aronin N, Chase K, Young C, et al. CAG expansion affects the expression of mutant Huntingtin in the Huntington’s disease brain. Neuron. Nov 1995;15(5):1193–201. doi:10.1016/0896-6273(95)90106-x

15. Iuliano M, Seeley C, Sapp E, et al. Disposition of Proteins and Lipids in Synaptic Membrane Compartments Is Altered in Q175/Q7 Huntington’s Disease Mouse Striatum. Front Synaptic Neurosci. 2021;13:618391. doi:10.3389/fnsyn.2021.618391

16. Sapp E, Valencia A, Li X, et al. Native mutant huntingtin in human brain: evidence for prevalence of full-length monomer. J Biol Chem. Apr 13 2012;287(16):13487–99. doi:10.1074/jbc.M111.286609

17. Vonsattel JP, Myers RH, Stevens TJ, Ferrante RJ, Bird ED, Richardson EP, Jr. Neuropathological classification of Huntington’s disease. J Neuropathol Exp Neurol. Nov 1985;44(6):559–77. doi:10.1097/00005072-198511000-00003

18. Alterman JF, Godinho B, Hassler MR, et al. A divalent siRNA chemical scaffold for potent and sustained modulation of gene expression throughout the central nervous system. Nat Biotechnol. Aug 2019;37(8):884–894. doi:10.1038/s41587-019-0205-0

19. O’Reilly D, Belgrad J, Ferguson C, et al. Di-valent siRNA-mediated silencing of MSH3 blocks somatic repeat expansion in mouse models of Huntington’s disease. Mol Ther. Jun 7 2023;31(6):1661–1674. doi:10.1016/j.ymthe.2023.05.006

20. Sapp E, Seeley C, Iuliano M, et al. Protein changes in synaptosomes of Huntington’s disease knock-in mice are dependent on age and brain region. Neurobiol Dis. Jul 2020;141:104950. doi:10.1016/j.nbd.2020.104950

21. Yamada K, Hariharan VN, Caiazzi J, et al. Enhancing siRNA efficacy in vivo with extended nucleic acid backbones. Nat Biotechnol. Aug 1 2024;doi:10.1038/s41587-024-02336-7

22. Narayanan KL, Chopra V, Rosas HD, Malarick K, Hersch S. Rho Kinase Pathway Alterations in the Brain and Leukocytes in Huntington’s Disease. Mol Neurobiol. May 2016;53(4):2132–40. doi:10.1007/s12035-015-9147-9

23. Kegel KB, Kim M, Sapp E, et al. Huntingtin expression stimulates endosomal-lysosomal activity, endosome tubulation, and autophagy. J Neurosci. Oct 1 2000;20(19):7268–78. doi:10.1523/JNEUROSCI.20-19-07268.2000

24. Kim YJ, Sapp E, Cuiffo BG, et al. Lysosomal proteases are involved in generation of N-terminal huntingtin fragments. Neurobiol Dis. May 2006;22(2):346–56. doi:10.1016/j.nbd.2005.11.012

25. Kegel KB, Sapp E, Alexander J, et al. Huntingtin cleavage product A forms in neurons and is reduced by gamma-secretase inhibitors. Mol Neurodegener. Dec 14 2010;5:58. doi:10.1186/1750-1326-5-58

26. DiFiglia M, Sapp E, Chase K, et al. Huntingtin is a cytoplasmic protein associated with vesicles in human and rat brain neurons. Neuron. May 1995;14(5):1075–81. doi:10.1016/0896-6273(95)90346-1

27. Deng Y, Wang H, Joni M, Sekhri R, Reiner A. Progression of basal ganglia pathology in heterozygous Q175 knock-in Huntington’s disease mice. J Comp Neurol. May 1 2021;529(7):1327–1371. doi:10.1002/cne.25023

28. Mattis VB, Tom C, Akimov S, et al. HD iPSC-derived neural progenitors accumulate in culture and are susceptible to BDNF withdrawal due to glutamate toxicity. Hum Mol Genet. Jun 1 2015;24(11):3257–71. doi:10.1093/hmg/ddv080

29. Shi Y, Kirwan P, Smith J, Robinson HP, Livesey FJ. Human cerebral cortex development from pluripotent stem cells to functional excitatory synapses. Nat Neurosci. Feb 5 2012;15(3):477–86, S1. doi:10.1038/nn.3041

30. Tousley A, Iuliano M, Weisman E, et al. Rac1 Activity Is Modulated by Huntingtin and Dysregulated in Models of Huntington’s Disease. J Huntingtons Dis. 2019;8(1):53–69. doi:10.3233/JHD-180311

31. Dragileva E, Hendricks A, Teed A, et al. Intergenerational and striatal CAG repeat instability in Huntington’s disease knock-in mice involve different DNA repair genes. Neurobiol Dis. Jan 2009;33(1):37–47. doi:10.1016/j.nbd.2008.09.014

32. Legleiter J, Lotz GP, Miller J, et al. Monoclonal antibodies recognize distinct conformational epitopes formed by polyglutamine in a mutant huntingtin fragment. J Biol Chem. Aug 7 2009;284(32):21647–58. doi:10.1074/jbc.M109.016923

33. Franich NR, Hickey MA, Zhu C, et al. Phenotype onset in Huntington’s disease knock-in mice is correlated with the incomplete splicing of the mutant huntingtin gene. J Neurosci Res. Dec 2019;97(12):1590–1605. doi:10.1002/jnr.24493

34. Hoschek F, Natan J, Wagner M, et al. Huntingtin HTT1a is generated in a CAG repeat-length-dependent manner in human tissues. Mol Med. Mar 8 2024;30(1):36. doi:10.1186/s10020-024-00801-2

35. Wheeler VC, Gutekunst CA, Vrbanac V, et al. Early phenotypes that presage late-onset neurodegenerative disease allow testing of modifiers in Hdh CAG knock-in mice. Hum Mol Genet. Mar 15 2002;11(6):633–40. doi:10.1093/hmg/11.6.633

36. Hickey MA, Zhu C, Medvedeva V, et al. Improvement of neuropathology and transcriptional deficits in CAG 140 knock-in mice supports a beneficial effect of dietary curcumin in Huntington’s disease. Mol Neurodegener. Apr 4 2012;7:12. doi:10.1186/1750-1326-7-12

37. Wheeler VC, White JK, Gutekunst CA, et al. Long glutamine tracts cause nuclear localization of a novel form of huntingtin in medium spiny striatal neurons in HdhQ92 and HdhQ111 knock-in mice. Hum Mol Genet. Mar 1 2000;9(4):503–13. doi:10.1093/hmg/9.4.503

38. Carroll JB, Lerch JP, Franciosi S, et al. Natural history of disease in the YAC128 mouse reveals a discrete signature of pathology in Huntington disease. Neurobiol Dis. Jul 2011;43(1):257–65. doi:10.1016/j.nbd.2011.03.018

39. Van Raamsdonk JM, Murphy Z, Slow EJ, Leavitt BR, Hayden MR. Selective degeneration and nuclear localization of mutant huntingtin in the YAC128 mouse model of Huntington disease. Hum Mol Genet. Dec 15 2005;14(24):3823–35. doi:10.1093/hmg/ddi407

40. Scherzinger E, Lurz R, Turmaine M, et al. Huntingtin-encoded polyglutamine expansions form amyloid-like protein aggregates in vitro and in vivo. Cell. Aug 8 1997;90(3):549–58. doi:10.1016/s0092-8674(00)80514-0

41. DiFiglia M, Sapp E, Chase KO, et al. Aggregation of huntingtin in neuronal intranuclear inclusions and dystrophic neurites in brain. Science. Sep 26 1997;277(5334):1990–3. doi:10.1126/science.277.5334.1990

42. Aldous SG, Smith EJ, Landles C, et al. A CAG repeat threshold for therapeutics targeting somatic instability in Huntington’s disease. Brain. May 3 2024;147(5):1784–1798. doi:10.1093/brain/awae063

43. Driscoll R, Hampton L, Abraham NA, et al. Dose-dependent reduction of somatic expansions but not Htt aggregates by di-valent siRNA-mediated silencing of MSH3 in HdhQ111 mice. Sci Rep. Jan 24 2024;14(1):2061. doi:10.1038/s41598-024-52667-3

44. Shelbourne PF, Keller-McGandy C, Bi WL, et al. Triplet repeat mutation length gains correlate with cell-type specific vulnerability in Huntington disease brain. Hum Mol Genet. May 15 2007;16(10):1133–42. doi:10.1093/hmg/ddm054

45. Vonsattel JP, DiFiglia M. Huntington disease. J Neuropathol Exp Neurol. May 1998;57(5):369–84. doi:10.1097/00005072-199805000-00001

46. Handsaker RE, Kashin S, Reed NM, et al. Long somatic DNA-repeat expansion drives neurodegeneration in Huntington disease. bioRxiv. 2024;doi:10.1101/2024.05.17.592722

47. Pressl C, Matlik K, Kus L, et al. Selective vulnerability of layer 5a corticostriatal neurons in Huntington’s disease. Neuron. Mar 20 2024;112(6):924–941 e10. doi:10.1016/j.neuron.2023.12.009

