## Supplemental figures for "Mutant Huntingtin exon1 protein detected in mouse brain with neoepitope antibody: effects of CAG repeat expansion, MSH3 silencing, and aggregation"

**Supplementary materials**

**Supplementary Table 1.** Control and Huntington’s disease brains used for western blot analysis of HTT1a.

|  | CAG | Category* | Age | Sex | Brain region | Source  (see footnote) | Other name | Publication |
| --- | --- | --- | --- | --- | --- | --- | --- | --- |
| 1060 | 17/17 | Control | 55 | m | Cortex | 1 | C1 | Sapp et al., 2012, Aronin et al., 1995 |
| 1063 | 17/22 | Control | 93 | f | Cortex | 1 | C2 | Aronin et al., 1995 |
| 1070 | 9/17 | Control | 68 | m | Cortex | 1 | C8 | Sapp et al., 2012, Aronin et al., 1995 |
| 206 | NA | Control | 49 | m | Cortex | 2 |  | Sapp et al., 2012 |
| A136 | NA | Control | NA | NA | Post. parietal cortex | 3 |  |  |
| Z021 | 12/17 | Control | 68 | f | Putamen | 1 |  |  |
| Z125 | 15/19 | Control | 64 | m | Cortex | 1 |  | Iuliano et al., 2021 |
| 1067 | 16/43 | HD3 | 62 | f | Cortex | 1 | A2 | Aronin et al., 1995 |
| 1069 | 17/43 | HD2 | 57 | m | Cortex | 1 | A3 | Sapp et al., 2012, Aronin et al., 1995 |
| 1065 | 25/40 | HD2 | 40 | f | Cortex | 1 | A11 | Sapp et al., 2012, Aronin et al., 1995 |
| 1429 | 18/41 | HD | 72 | m | Putamen | 4 |  | Iuliano et al., 2021 |
| 253 | 20/43 | HD1 | 45 | f | Cortex | 2 |  |  |
| 210 | 43/48 | HD | 43 | f | Cortex | 2 |  | Sapp et al., 2012 |
| A143 | NA | HD1 | NA | NA | Post. parietal cortex | 3 |  |  |
| A122 | NA | HD4 | NA | NA | Post. parietal cortex | 3 |  |  |
| Z076 | 18/53 | HD | 29 | m | Cortex | 1 |  |  |
| 3053 | 27/42 | HD3 | 66 | f | Cortex and putamen | 5 | A12 | Sapp et al., 2012, Aronin et al., 1995 |
| 1068 | 15/69 | HD3 | 28 | f | Cortex | 1 | J5 | Sapp et al., 2012, Aronin et al., 1995 |
| 1066 | 17/105 | HD4 | 12 | f | Cortex | 1 | J6 | Aronin et al., 1995 |
| 4501 | NA | HD4 | 8 | f | Cortex | 2 |  |  |
| Z067 | NA | PD | 84 | f | Cortex | 1 |  |  |
| Z069 | NA | PD | 77 | f | Cortex | 1 |  |  |
| Z078 | NA | PD | 31 | f | Cortex | 1 |  |  |

^1^ Massachusetts General Hospital Neuropharmacology Laboratory Brain Bank, ^2^ New York Brain Bank at Columbia University, ^3^ Anton Reiner, University of Tennessee, ^4^ Massachusetts Alzheimer's Disease Resource Center, ^5^ Harvard Brain Tissue Resource Center; *Number represents Huntington’s disease grade based on Vonsattel grading system when neuropathological classification was performed;^1^ NA=not available; HD=Huntington’s disease; PD=Parkinson’s disease; Post.=posterior.

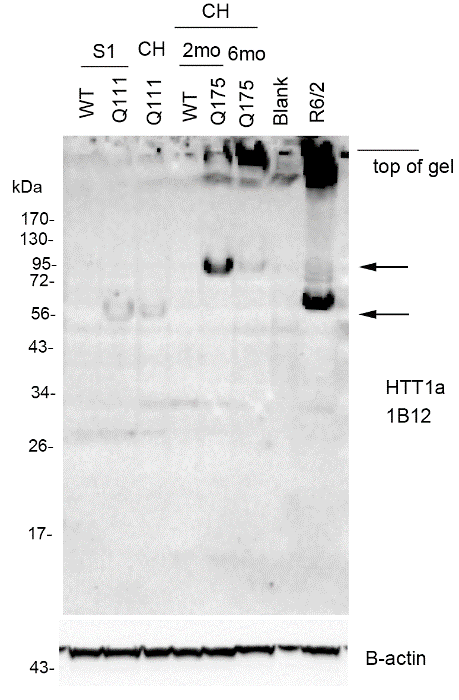

**Supplementary Fig. 1 HTT1a bands migrate faster in a 12% Bis-tris gel.** 20 µg samples from different preparations (S1 fraction and CH=crude homogenate) from R6/2, WT, Q111 and Q175 mice were separated using 12% Bis-tris 15-well gels. Arrows indicate the position of the HTT1a bands detected with 1B12 at about ~60 kDa in 12-week Q111 mice and at 90 kDa in Q175 mice that migrate faster and with less signal intensity than in 3-8% Tris-acetate gels (see **Figs. 1-4** and **Table 1**). The HTT1a smear in Q175 at 6 months and in the R6/2 mouse is prominent and more concentrated at the top of the blot. Note the changes in abundance of HTT1a band and smear between 2- and 6-month old Q175 mice as seen in the 3-8% Tris-acetate gels (**Fig 2C**). Mouse labels are positioned at the center of each lane. “Blank” marks a lane which had no protein loading but some protein spillover into the lane occurred. Full blots are shown in **Supplementary Fig. 8.**

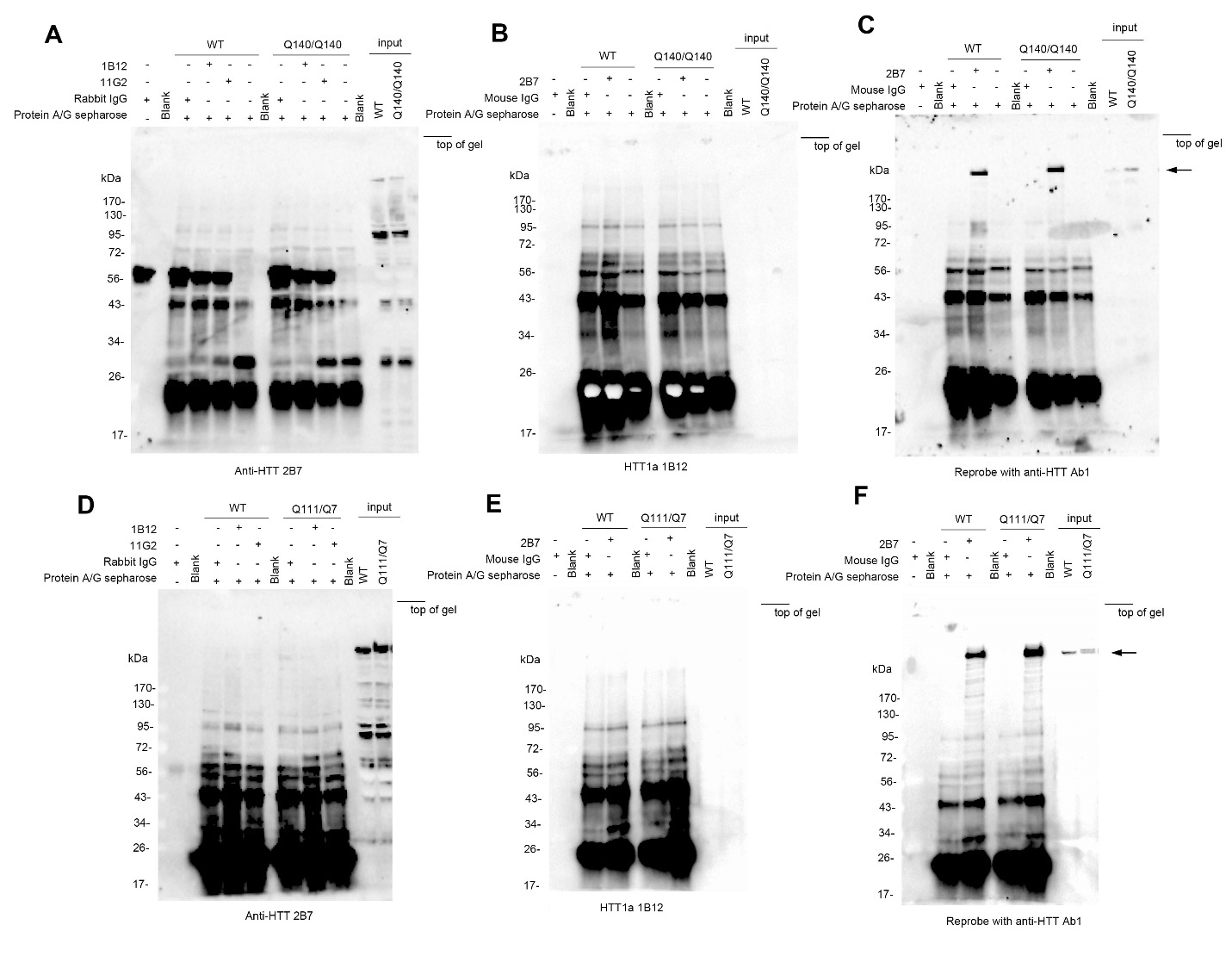

**Supplementary Fig. 2 Immunoprecipitation with antibodies 1B12, 11G2 and 2B7 in lysates from Q140/Q140 and Q111/Q7 mouse brain.** Immunoprecipitation assays were performed on WT and 6-month old Q140/Q140 (**A-C**) and 3-month old Q111/Q7 (**D-F**) mouse caudate putamen using 5 µg 1B12 and 11G2 antibodies and probing with anti-HTT antibody 2B7 (**A, D**) or 5 µg 2B7 antibody and probing with 1B12 (**B, E**) as described in methods. Under these conditions, there are no specific bands in the Huntington’s disease Q140/Q140 or Q111/Q7 caudate putamen immunoprecipitated with 1B12 and 11G2 antibodies. Full length HTT is present and enriched in immunoprecipitates obtained with 2B7 antibody and probed with antibody Ab1 (arrows in **C, F**). All immunoprecipitation samples were separated using 4-12% Bis-tris 26-well gels. At the top of the blots, labels are positioned at the center of each lane. “Blank” marks a lane with no protein loading. Monoclonal 2B7 and polyclonal Ab1 antibodies were made to HTT1-17.

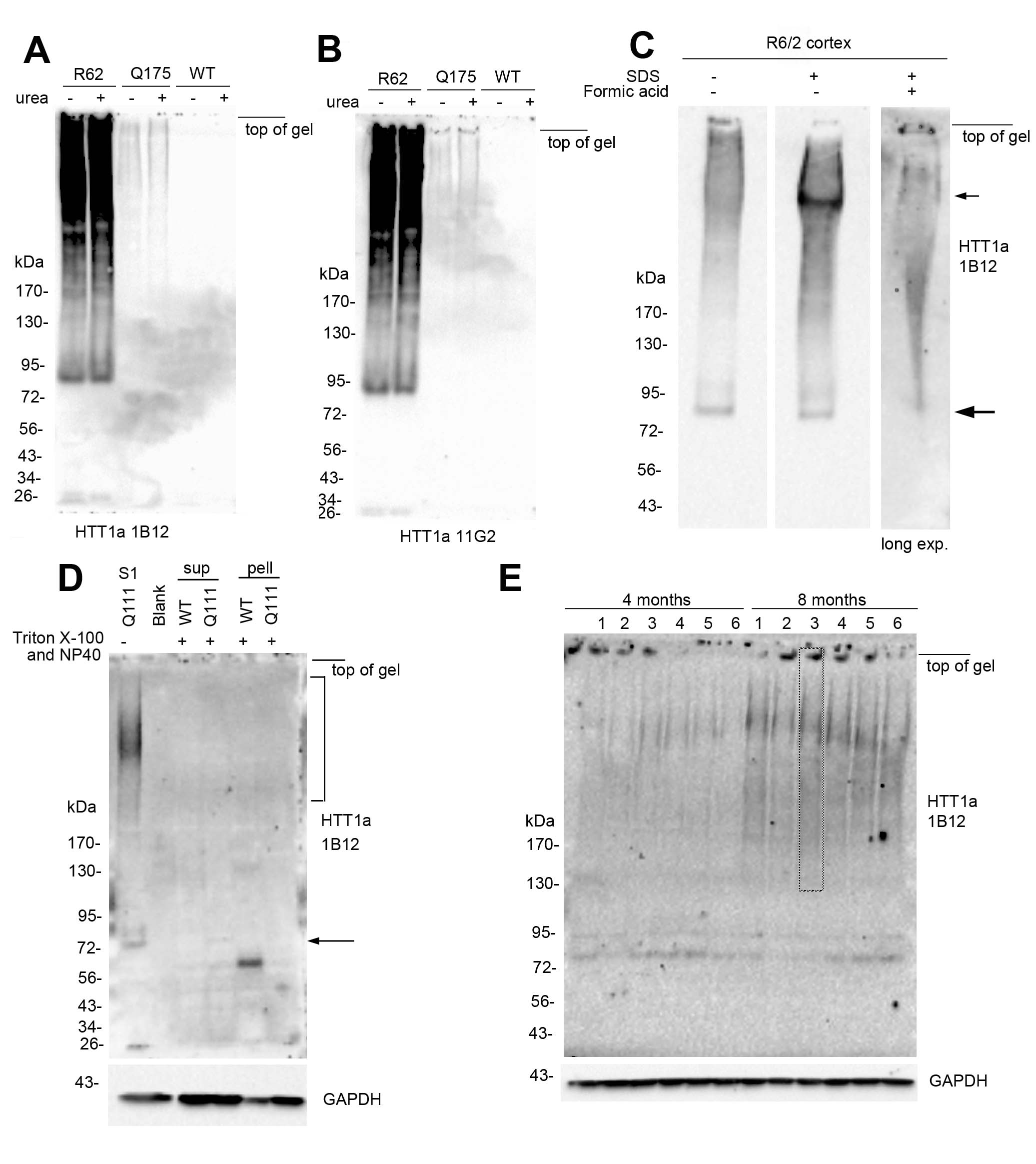

**Supplementary Fig. 3 A, B, C**. **Solubilizing crude homogenates with 8M urea or formic acid had minimal effect on detecting HTT1a.** Lysates from Huntington’s disease mice (R62, Q175) were incubated without (-) or with (+) 8M urea at room temperature for 30 minutes following a previously published method.^2^ There was no change with urea treatment in HTT1a signal detected with antibodies 1B12 (**A**) or 11G2 (**B**). **C**. Sequential treatment of crude homogenates from R6/2 cortex with SDS and formic acid was performed according to method described by Landles et al., 2010.^3^ Solubilization with SDS alone increased the high molecular mass (HMM) HTT1a smear and increased a HMM band (small arrow) but no change occurred in 85kDa HTT1a band (large arrow). Shorter exposures than those in **A** and **B** are shown so the distinct band at the small arrow is distinguishable. Addition of formic acid treatment resulted in the disappearance of the HMM band (small arrow) and clearance of the HTT1a band (large arrow). **D**. To test use of mild detergent, the method of Kim et al. (2006) was followed to extract protein from WT and 12 week old Q111 caudate putamen using 20mM Tris pH 7.2, 150mM NaCl, 1% Triton X-100 and 0.1% NP40.^4^ After 16000xg centrifugation, the soluble supernatants (sup) and detergent insoluble pellets (pell) showed minimal signal for the 72 kDa band (arrow) and no HMM smear (bracket) compared to the detergent-free supernatant (S1). **E**. Western blot of additional samples used for quantification of the YAC128 4- and 8- month old mice in the graph shown in **Fig 4D**. Crude homogenates from 4- and 8-month-old YAC128 mouse striatum were separated by SDS-PAGE and probed with antibody 1B12. Signal intensity for the HTT1a smear indicated by dashed lines in 8-month lane 3 was measured in the same area for each sample and normalized to GAPDH loading control. 20 μg samples were separated using 3-8% Tris-acetate gels for all panels. At the top of the blots, labels or mouse numbers are positioned at the center of each lane. Full blots are shown in **Supplementary Fig. 9.**

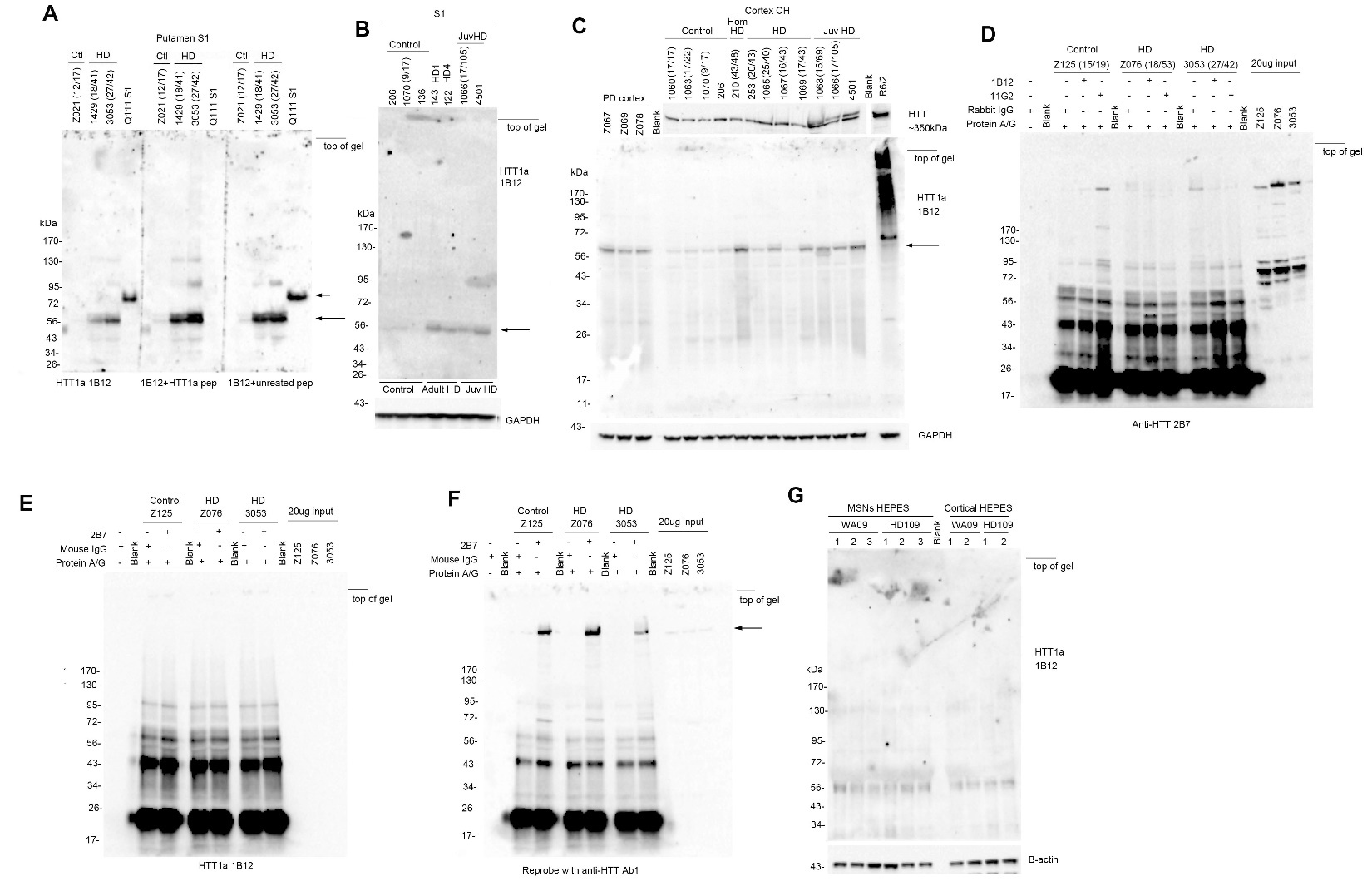

**Supplementary Fig. 4 Use of blocking peptide and immunoprecipitation assays show that 1B12 antibody does not detect HTT1a in human patient putamen or cortex. A**. Using a 3-8% Tris-acetate 15-well gel and 20 μg samples, a doublet band detected at 56-60 kDa (long arrow) in human Huntington’s disease putamen but not in control (Ctl) S1 fraction was not blocked by preincubation of antibody 1B12 with HTT1a peptide for 1 hour at room temperature. One Q111 S1 sample was run on the same gel as a positive control and the signal was blocked with the HTT1a peptide (short arrow, middle blot) but not with an unrelated peptide. **B**. There were no CAG-length dependent bands detected in the cortex S1 fraction of juvenile-onset Huntington’s disease patients (JuvHD), one with a known CAG repeat of 105. 20 μg samples were separated on a 3-8% Tris-acetate 26-well gel and the 56-60 kDa doublet is detected (long arrow). **C**. The signal for 56-60 kDa doublet was increased in Parkinson’s disease cortex crude homogenate (CH) as well as Huntington’s disease cortex compared to control samples (long arrow) when 20 μg samples were separated on a 26-well 12% Bis-tris gel. **D**. Immunoprecipitation with 1B12 or 11G2 from control (Z125) and Huntington’s disease (Z076 and 3053) post-mortem cortex and probed with N-terminal anti-HTT antibody 2B7 did not detect HTT1a bands specific to the Huntington’s disease samples above the background signal detected in the no antibody lane. **E, F.** Immunoprecipitation of HTT with 2B7 antibody from same the control and Huntington’s disease post-mortem cortex lysates as in **D** did not immunoprecipitate HTT1a when detected with antibody 1B12 (**E**) but detected and enriched for full length HTT after blots in **E** were stripped and re-probed with N-terminal anti-HTT antibody Ab1 (arrow in **F**). Immunoprecipitation samples were separated on 4-12% Bis-tris 26 well gels (**D-F**). **G**. The 56-60 kDa doublet was detected in control (WA09) and HD109 medium spiny neurons and cortical neurons with 1B12 antibody indicating it is not HD specific. Neuron samples (20 μg) were separated on a 3-8% Tris-acetate 26-well gel. At the top of the blots, labels are positioned at the center of each lane. “Blank” marks a lane without protein. Full blots are shown in **Supplementary Fig. 10**.

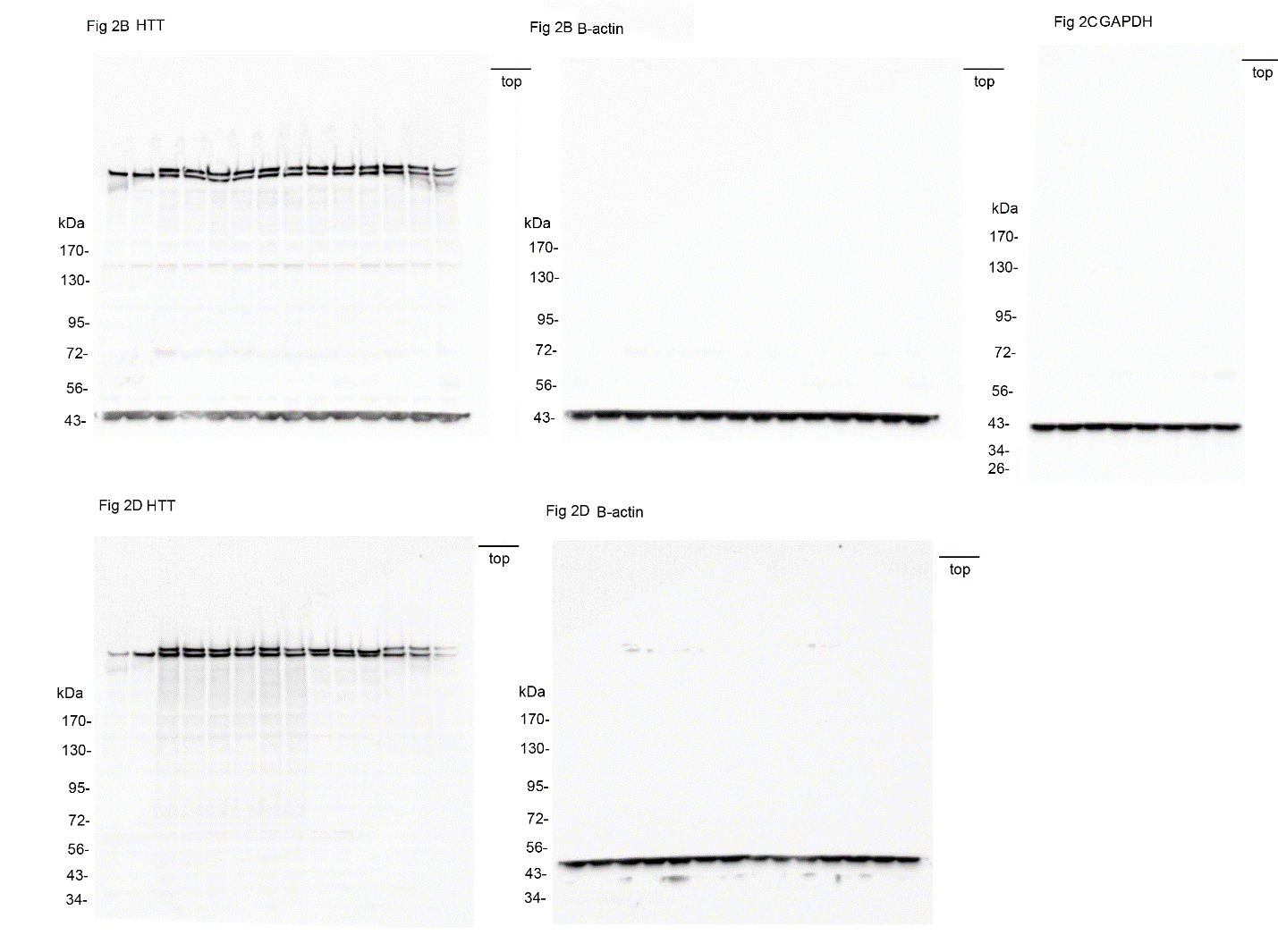

**Supplementary Fig. 5 Full blots shown in Fig. 2.** Please note that blots were reprobed with different antibodies and residual signal from the previous antibody may be visible

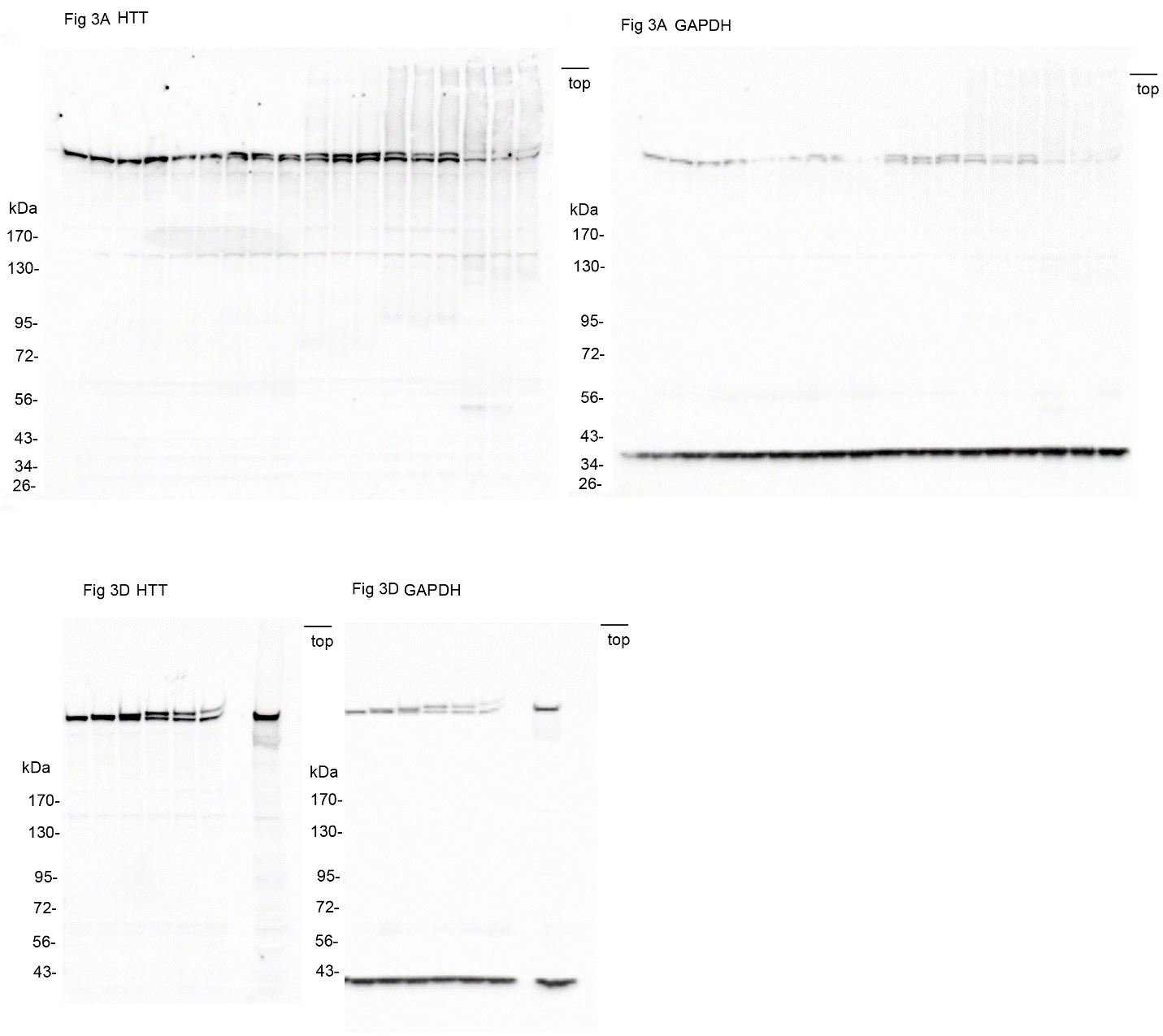

**Supplementary Fig. 6. Full blots shown in Fig. 3.** Please note that blots were reprobed with different antibodies and residual signal from the previous antibody may be visible.

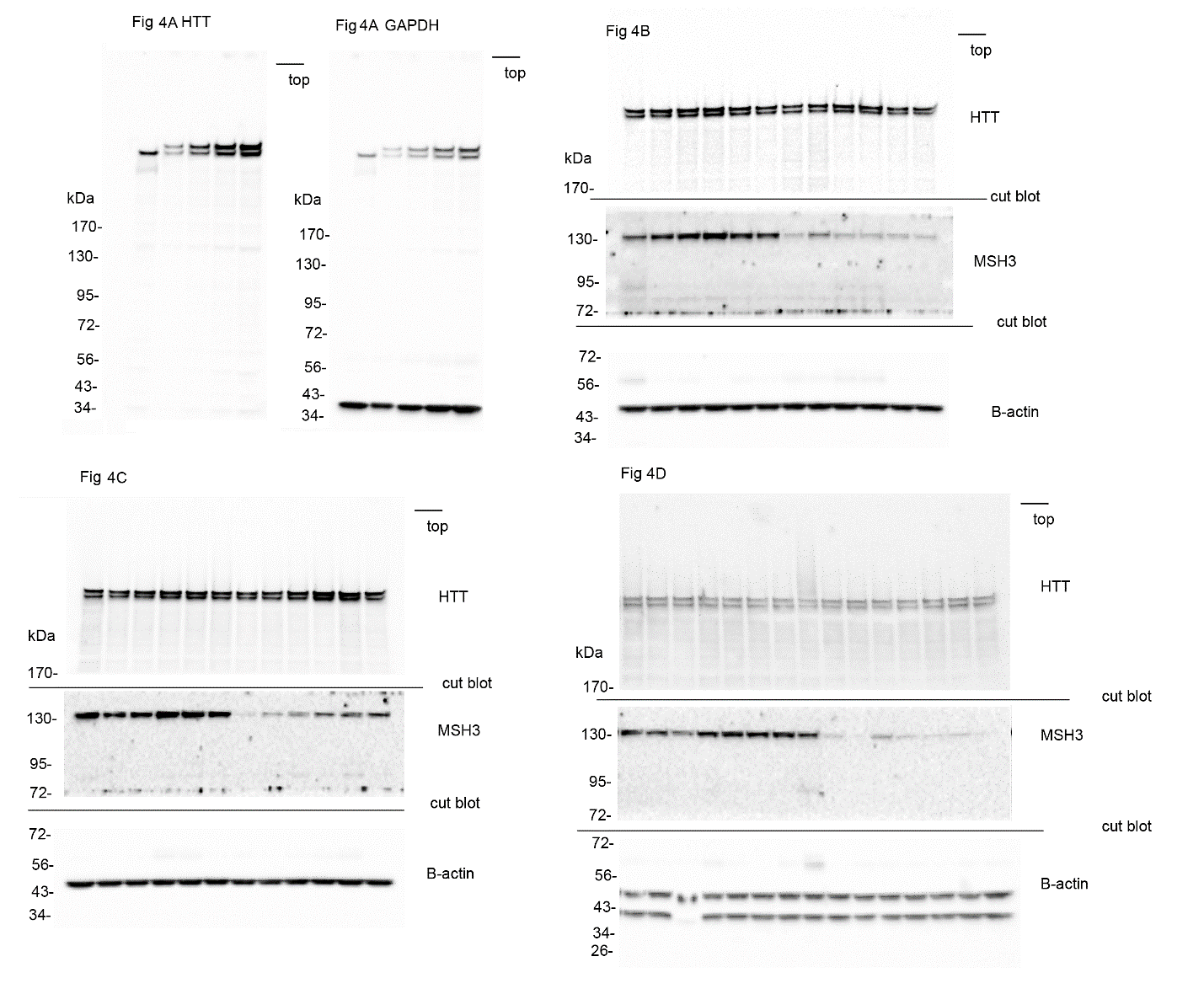

**Supplementary Fig. 7. Full blots shown in Fig. 4.** Please note that blots were reprobed with different antibodies and residual signal from the previous antibody may be visible. For Fig. 4B-D, blots were cut into strips and each strip was probed with a different antibody.

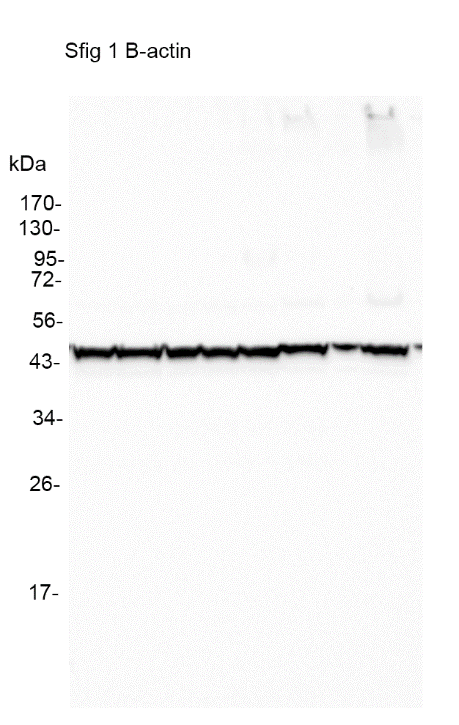

**Supplementary Fig. 8 Full blot shown in Supplementary Fig. 1.** Please note that blots were reprobed with different antibodies and residual signal from the previous antibody may be visible.

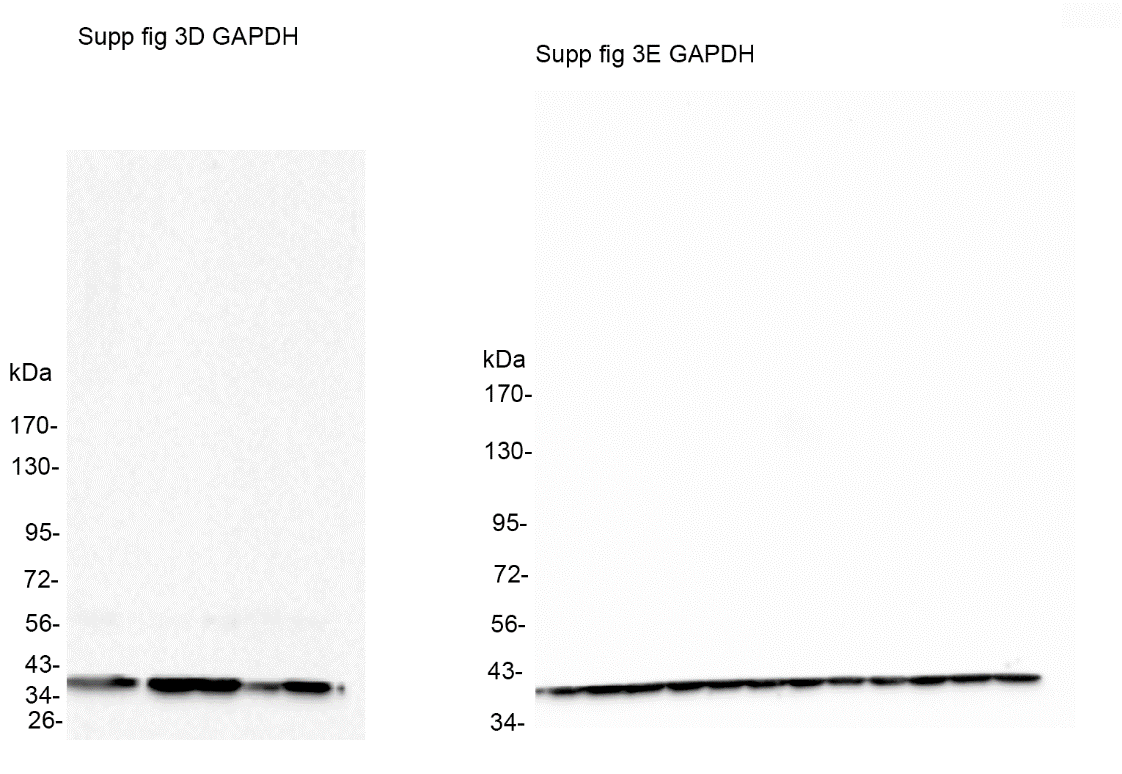

**Supplementary Fig. 9 Full blots shown in Supplementary Fig. 3.**

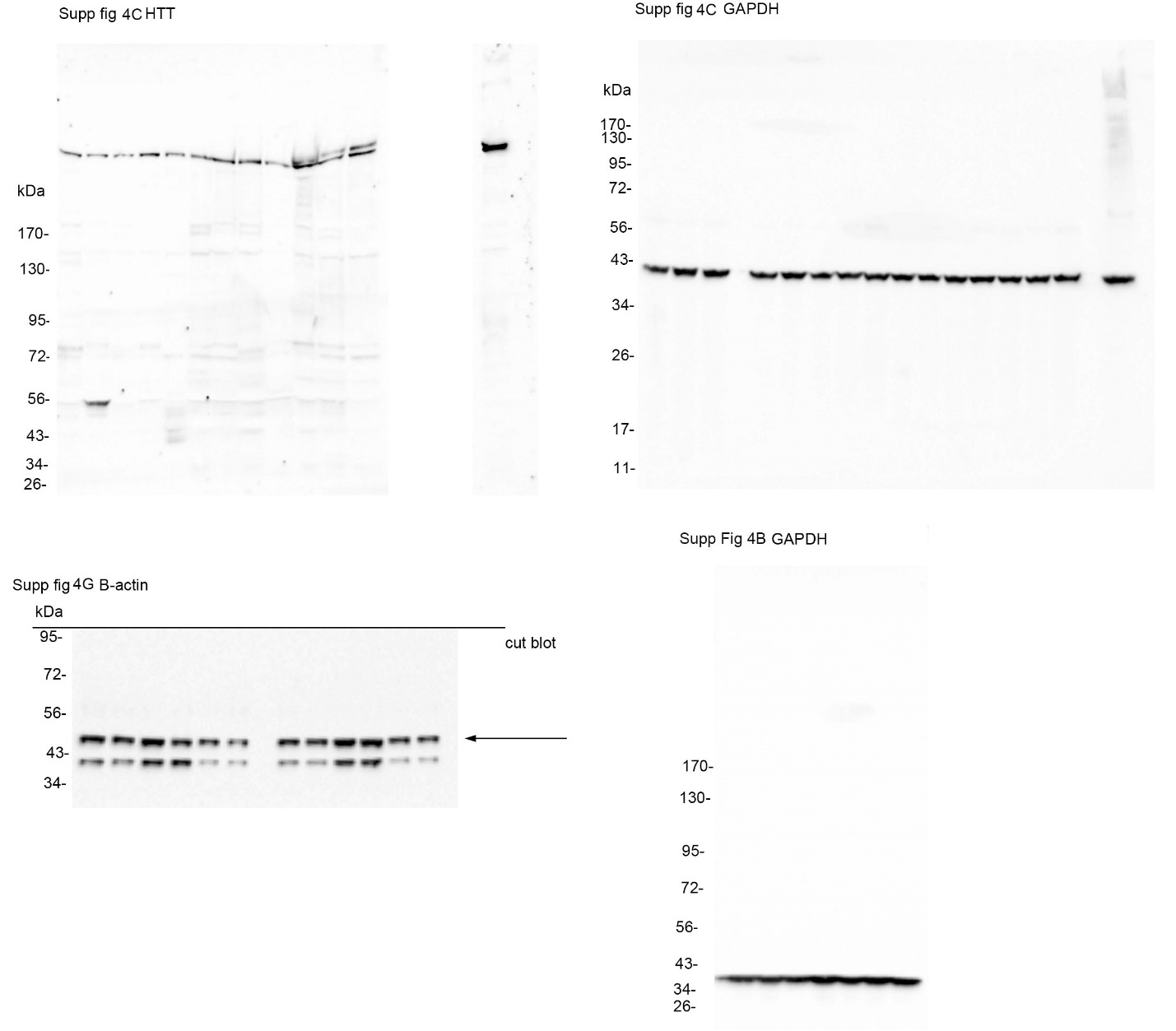

**Supplementary Fig. 10 Full blots shown in Supplementary Fig. 4.** Please note that blots were reprobed with different antibodies and residual signal from the previous antibody may be visible. For Supplementary Fig. 4G blot was cut into strips and before probing with different antibodies.

**Supplementary References**

1.	Vonsattel JP, Myers RH, Stevens TJ, Ferrante RJ, Bird ED, Richardson EP, Jr. Neuropathological classification of Huntington's disease. *J Neuropathol Exp Neurol*. Nov 1985;44(6):559-77. doi:10.1097/00005072-198511000-00003

2.	Sapp E, Valencia A, Li X, et al. Native mutant huntingtin in human brain: evidence for prevalence of full-length monomer. *J Biol Chem*. Apr 13 2012;287(16):13487-99. doi:10.1074/jbc.M111.286609

3.	Landles C, Sathasivam K, Weiss A, et al. Proteolysis of mutant huntingtin produces an exon 1 fragment that accumulates as an aggregated protein in neuronal nuclei in Huntington disease. *J Biol Chem*. Mar 19 2010;285(12):8808-23. doi:10.1074/jbc.M109.075028

4.	Kim YJ, Sapp E, Cuiffo BG, et al. Lysosomal proteases are involved in generation of N-terminal huntingtin fragments. *Neurobiol Dis*. May 2006;22(2):346-56. doi:10.1016/j.nbd.2005.11.012
